# The structural basis for RNA slicing by human Argonaute2

**DOI:** 10.1101/2024.08.19.608718

**Authors:** Abdallah A. Mohamed, Peter Y. Wang, David P. Bartel, Seychelle M. Vos

**Affiliations:** Department of Biology, Massachusetts Institute of Technology, 31 Ames Street, Cambridge, MA, 02139, USA; Whitehead Institute for Biomedical Research, 455 Main Street, Cambridge, MA, 02142, USA; Howard Hughes Medical Institute, Cambridge, MA, 02142, USA

## Abstract

Argonaute (AGO) proteins associate with guide RNAs to form complexes that slice transcripts that pair to the guide. This slicing drives post-transcriptional gene-silencing pathways that are essential for many eukaryotes and the basis for new clinical therapies. Despite this importance, structural information on eukaryotic AGOs in a fully paired, slicing-competent conformation—hypothesized to be intrinsically unstable—has been lacking. Here we present the cryogenic-electron microscopy structure of a human AGO−guide complex bound to a fully paired target, revealing structural rearrangements that enable this conformation. Critically, the N domain of AGO rotates to allow the RNA full access to the central channel and forms contacts that license rapid slicing. Moreover, a conserved loop in the PIWI domain secures the RNA near the active site to enhance slicing rate and specificity. These results explain how AGO accommodates targets possessing the pairing specificity typically observed in biological and clinical slicing substrates.

## INTRODUCTION

The AGO protein family is found in all domains of life.^1^ In eukaryotes, AGOs typically associate with ∼22-nt guide RNAs, including microRNAs (miRNAs) and endogenous small interfering RNAs (siRNAs), to form an RNA-induced silencing complex (RISC), which slices RNA transcripts that pair extensively to the guide.^2,3^ This slicing activity is central to RNA interference (RNAi),^3^ a gene-silencing pathway critical for defense against viruses and transposons.^4,5^ Slicing activity also regulates endogenous cellular transcripts^6–11^ and has been harnessed for mRNA knockdown—both as a research tool^12^ and for clinically approved therapies^13^.

Among the four human AGO paralogs (HsAGO1, -2, -3, and -4), HsAGO2 has retained the ancestral endonuclease activity that slices extensively complementary RNAs.^14,15^ As is typical of AGO proteins, HsAGO2 has four major structural domains (N, PAZ, MID, and PIWI) connected by two linker domains (L1 and L2), together creating a Y-shaped RNA-binding channel bifurcated by the N domain (**Figure 1A**).^16,17^ The guide RNA also has four regions, known as the seed (nucleotides 2–8), central (nucleotides 9–12), supplementary (nucleotides 13–16), and tail (nucleotides 17–22) regions.

**Figure 1.**
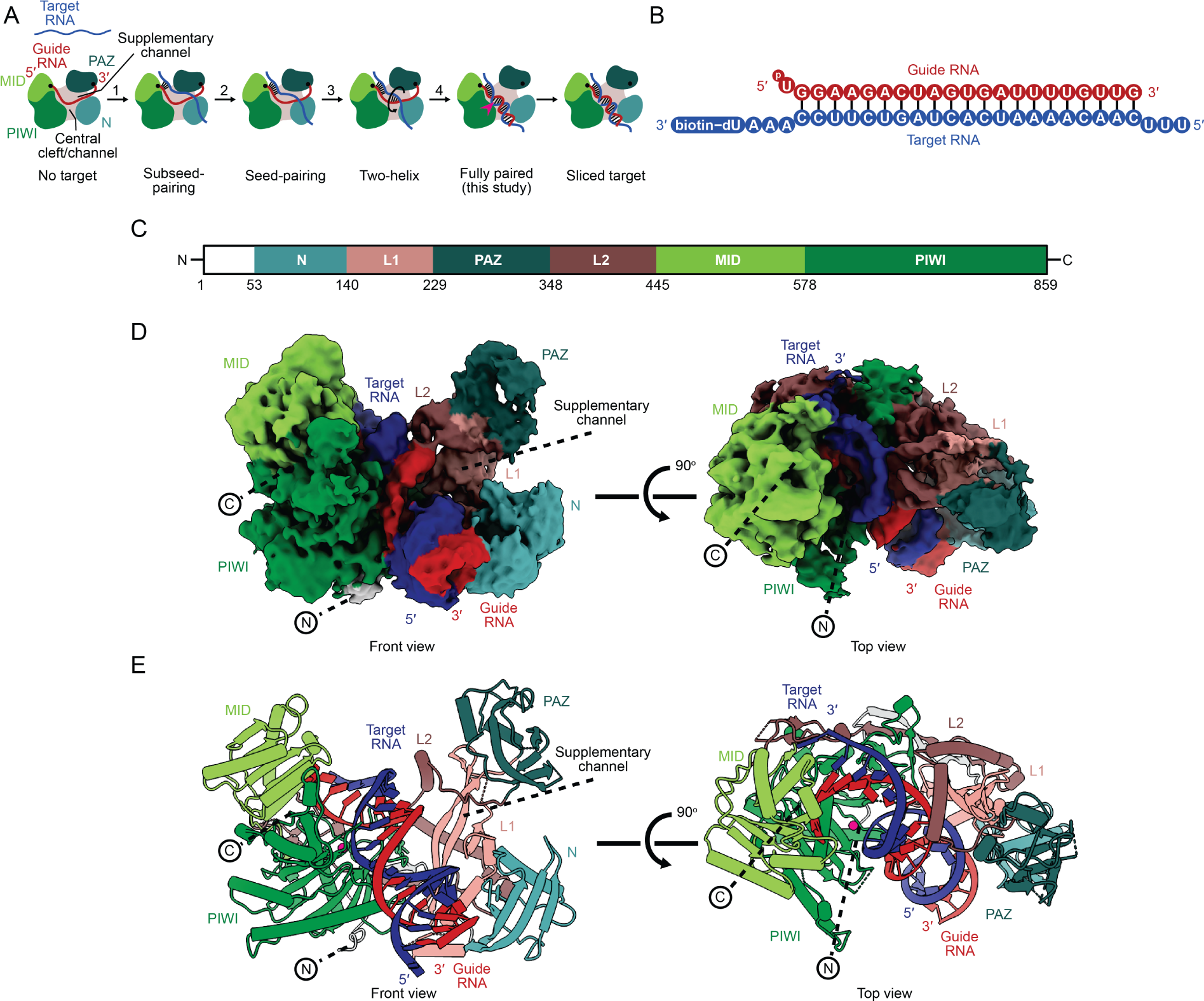
Structure of the HsAGO2−miR-7−target complex in the fully paired conformation. (A) Four-step model for the RNA conformational changes required to achieve the fully paired conformation. Guide and target RNAs are colored red and blue, respectively, and the MID, PIWI, PAZ, and N domains are colored light green, dark green, teal, and cyan, respectively. RNA-binding channels are labeled. A pink caret indicates the active site. (B) Schematic of the guide−target RNA duplex used for structural analysis. Vertical black lines represent base pairing. Biotin attachment to a deoxyuridine nucleotide is indicated. (C) Primary structure of HsAGO2 with residue numbers indicating domain boundaries. (D) Cryo-EM structure of the HsAGO2−miR-7−target complex in a fully paired conformation. Map (Map 2) used for modeling is shown from front and top views. RNAs are colored as in panel **B**. HsAGO2 is colored as in panel **C**. (E) Cryo-EM structure of the HsAGO2−miR-7−target complex in a fully paired conformation. Model is shown from front and top views. RNAs are colored as in panel **B**. HsAGO2 is colored as in panel **C**.

Pairing of the guide to a fully complementary target RNA poses conformational challenges for AGO RISC of humans and other animals.^18^ For example, contacts between the guide and protein impede the rotational movement required for guide and target strands to wrap around each other. Moreover, a narrow central cleft within the protein hinders propagation of pairing through the central region of the guide.^19^ A four-step pathway, supported by structural studies, is proposed to overcome these challenges (**Figure 1A**).^2^ In step one, pairing to target RNA initiates at guide-RNA positions 2−5, which are pre-organized by the MID and PIWI domains in an A-form-like conformation poised to nucleate initial pairing.^20–26^ In step two, pairing propagates to the remainder of the seed region, which is accommodated by repositioning of helix-7 of the protein;^19,21,22,27^ position 1 is not base-paired but is instead recognized by the HsAGO2 protein, which favors an adenosine at this position.^28–30^ In step three, a second helix nucleates in a widened supplementary channel, at or near guide-RNA positions 13−16,^31^ where pairing can resume without the target strand having to pass between the guide and the protein. Finally, in step four, the second helix rolls from the supplementary channel, over the N domain, into the central channel, in concert with propagation of pairing at its two flanks—both towards the guide 3′ terminus and within the central region.^2^ The multiple conformational changes occurring during this last step each appear to enable or promote the others.^32^

Pairing to the central region brings the target scissile phosphate into the active site, where it can be sliced. However, a structure with this centrally paired target has not been determined for a metazoan RISC. An attempt to crystalize HsAGO2 in a slicing-competent conformation has yielded only a two-helix conformation stalled at step three, with the target scissile phosphate positioned far away (>10 Å) from the active site (**Figure S1A**).^31^ Thus, the structural framework of the metazoan slicing-competent conformation is primarily inferred by analogy to the structures of a prokaryotic RISC (TtAGO from *Thermus thermophilus*) (**Figure S1B**) and a plant RISC (AtAGO10 from *Arabidopsis thaliana*) with centrally paired targets, obtained by X-ray crystallography and cryogenic-electron microscopy (cryo-EM), respectively.^33–35^ Meanwhile, biophysical exploration of the slicing-competent conformation of HsAGO2 comes from lower-resolution studies, including single-molecule fluorescence-resonance-energy-transfer (FRET) microscopy and chemical footprinting.^32,36^

Perhaps even more consequential than the absence of a metazoan slicing structure, is the absence of a eukaryotic slicing structure with pairing beyond position 16. Of particular interest is the question of how a contiguously paired, guide–target duplex is accommodated. The lone eukaryotic RISC structure showing a centrally paired, slicing-competent conformation is of AtAGO10, but in this structure pairing does not extend beyond guide nucleotide 16.^34^ Moreover, when pairing is modeled to extend to the end of the guide, it clashes with the N domain, implying that to accommodate additional pairing to the guide RNA tail, either the N domain must move or be remodeled, or the duplex must bend. Precedents for both N-domain movement and duplex bending are found in recent structures of fully paired bacterial RISCs^37,38^. Alternatively, bending can be achieved through incomplete pairing of the central region^31,34^ (**Figures S1C and S1D**), and by analogy to PIWI, an AGO paralog, accommodating pairing beyond position 16 might drive opening of the central cleft to the point that the N−PAZ lobe is no longer structurally coupled to the remainder of the complex.^39^

The AtAGO10 slicing structure lacks pairing beyond position 16 because it was determined using a target that ends at position 16, with the idea that this truncated target would prevent guide-strand unloading while favoring the centrally paired, slicing-competent conformation.^34^ This idea is supported by two lines of evidence. First, in the structure that initially revealed RISC in a centrally paired, slicing-competent conformation, i.e., a structure of TtAGO RISC with a complementary target, the N domain blocks pairing beyond position 16, even though the target has potential to pair up to guide position 19.^33^ This observation led to the idea that disrupting pairing to the tail region of the guide might be required to achieve the slicing conformation. Second, biochemical analyses with a focus on two guide RNAs (let-7a and miR-21) indicate that pairing beyond position 16 can be dispensable or even somewhat detrimental for efficient slicing.^40,41^ Countering these two lines of evidence are results of more recent structural and biochemical studies. Structural analyses of other prokaryotic AGOs have revealed MpAGO (from *Marinitoga piezophila*) in a slicing conformation with pairing that extends up to position 20 (**Figure S1E**).^38^ Moreover, recent analyses of HsAGO2 with a broader range of guide RNAs shows that for most, and particularly those with AU-rich central regions, pairing beyond position 16 is not detrimental for slicing but instead required for efficient slicing, and chemical-footprinting results support the facile formation and high occupancy of the fully paired conformation, regardless of the guide-RNA sequence.^32^ These more recent results concur with the observation that known slicing sites of miRNAs and endogenous siRNAs of animals and plants, as well as target sites of synthetic siRNAs in the clinic, typically pair extensively throughout the length of the guide—not to just nucleotides 2–16.^6–10,13^

Studies of the N domain also hint at a function for base pairing to the end of the guide. Domain-swaps of HsAGO paralogs show that the N domain of HsAGO2 is required to fully restore slicing activity to noncatalytic paralogs, suggesting that the N domain, which abuts pairing at the tail region, harbors features that enable slicing.^42–44^ Furthermore, deleting the N–PAZ lobe results in more permissive slicing, supporting a model in which the N−PAZ lobe governs slicing through steric effects.^45,46^ The mechanisms by which the N domain contributes to slicing, however, await information on how this domain accommodates and contacts the fully paired guide.

To determine the slicing-competent conformation of a metazoan RISC and to learn how RISC can accommodate and contact a fully paired target RNA, we set out to use single-particle cryo-EM to determine the structure of miR-7–HsAGO2 RISC fully paired to its target.

## RESULTS

### Structure of HsAGO2 in the slicing conformation

FLAG-tagged HsAGO2 harboring a catalytically dead mutation (D669A) was expressed in human cells and loaded with a 22-nt miR-7 miRNA in cell lysate. The resulting miR-7−HsAGO2 RISC was enriched based on FLAG affinity, incubated with a biotinylated target RNA (**Figure 1B**), then purified with target using Strep-Tactin affinity, followed by size-exclusion chromatography (**Figures S1F and S1G**).

We found that samples applied to grids with a thin carbon support film produced mono-disperse, single particles, which we analyzed by cryo-EM. After extensive classification, we obtained a map with good occupancy for a continuous RNA duplex, as well as the MID and PIWI domains of HsAGO2, and lower occupancy for the N and PAZ domains, presumably due to their greater flexibility. To better resolve the N and PAZ domains, we performed 3D classification with a focus mask along with subsequent classification and refinements and obtained a map with increased occupancy of the N and PAZ domains (**Figure S2**). To overcome modest orientation bias in this map, we employed spIsoNet.^47^ The final map had a nominal resolution of 3.3 Å (gold standard FSC 0.143) (**Figures 1C–1E and S3, Movie S1**), with the active site achieving a resolution of 3.3 Å and the N and PAZ domains ranging from 5–8 Å in resolution (**Figure S3C**). Modeling of the protein domains was guided by an X-ray crystal structure of HsAGO2 without bound target.^24^ Modeling of RNA nucleotides was guided by either the structure of HsAGO2 in a two-helix conformation^31^ (positions 1–8) or an A-form RNA duplex (positions 9–22).

The model was manually adjusted, real-space refined, and shows good stereochemistry (**Table 1**).

**Table 1.** Cryo-EM data collection, refinement, and validation statistics.

|  | HsAGO2 slicing-competent complex<br>(EMD-45752)<br>(PDB: 9CMP) |
| --- | --- |
| **Data collection and processing** |  |
| Magnification | 130,000 |
| Voltage (kV) | 300 |
| Electron exposure (e<sup>-</sup>/Å<sup>2</sup>) | 51.0 |
| Defocus range (μm) | 0.2-1.4 |
| Pixel size (Å) |  |
| Symmetry imposed | C1 |
| Initial particle images (no.) | 1,427,327 |
| Final particle images (no.) | 69,008 |
| Map resolution (Å) | 3.3 |
| FSC threshold | 0.143 |
| Map resolution range (Å) | 3.3-8.0 |
| **Refinement** |  |
| Initial models used (PDB code) | 4OLA, 6N4O |
| Map sharpening *B* factor (Å<sup>2</sup>) | -100 |
| **Model composition** |  |
| Non-hydrogen atoms | 6218 |
| Protein residues | 768 |
| Nucleotides | 43 |
| Ligands | MG: 1 |
| ***B* factors (Å<sup>2</sup>)** |  |
| Protein | 128.41 |
| Nucleotide | 130.03 |
| Ligand | 70.30 |
| **R.m.s. deviations** |  |
| Bond lengths (Å) | 0.003 |
| Bond angles (°) | 0.657 |
| **Validation** |  |
| MolProbity score | 1.15 |
| Clashscore | 1.43 |
| Poor rotamers (%) | 0.46 |
| **Ramachandran plot** |  |
| Favored (%) | 96.08 |
| Allowed (%) | 3.92 |
| Disallowed (%) | 0.00 |

### The N domain moves to accommodate the fully paired RNA duplex

Our map revealed the centrally paired, slicing-competent conformation of a metazoan RISC, thereby providing the final snapshot for the four-step pathway of target association (**Figure 1A**). This structure, together with structures of conformations analogous to the preceding intermediate states,^19,30,31,48^ enabled visualization of metazoan RISC traversing the four steps of target association (**Movie S2**).

Density for the RNA showed that the guide–target duplex was fully paired from position 2 to the end of the guide RNA, with distortions from A-form helix no larger than those observed for a 2−16-paired duplex within AtAGO10.^34^ This unimpeded path of the RNA duplex implied substantial movement or remodeling of the N domain. Indeed, comparison of our model with that of the two-helix HsAGO2 structure^31^ showed that the N domain moves dramatically to accommodate and potentially contact positions 15−19 of the guide–target duplex (**Figure 2A**). Specifically, the first alpha helix of the protein (α1) moves by 9 Å to widen the central channel (**Figure 2A**). Furthermore, β3−β7 move by ∼10−12 Å, together with the remainder of the N domain, to make room for the RNA duplex in the central channel.

**Figure 2.**
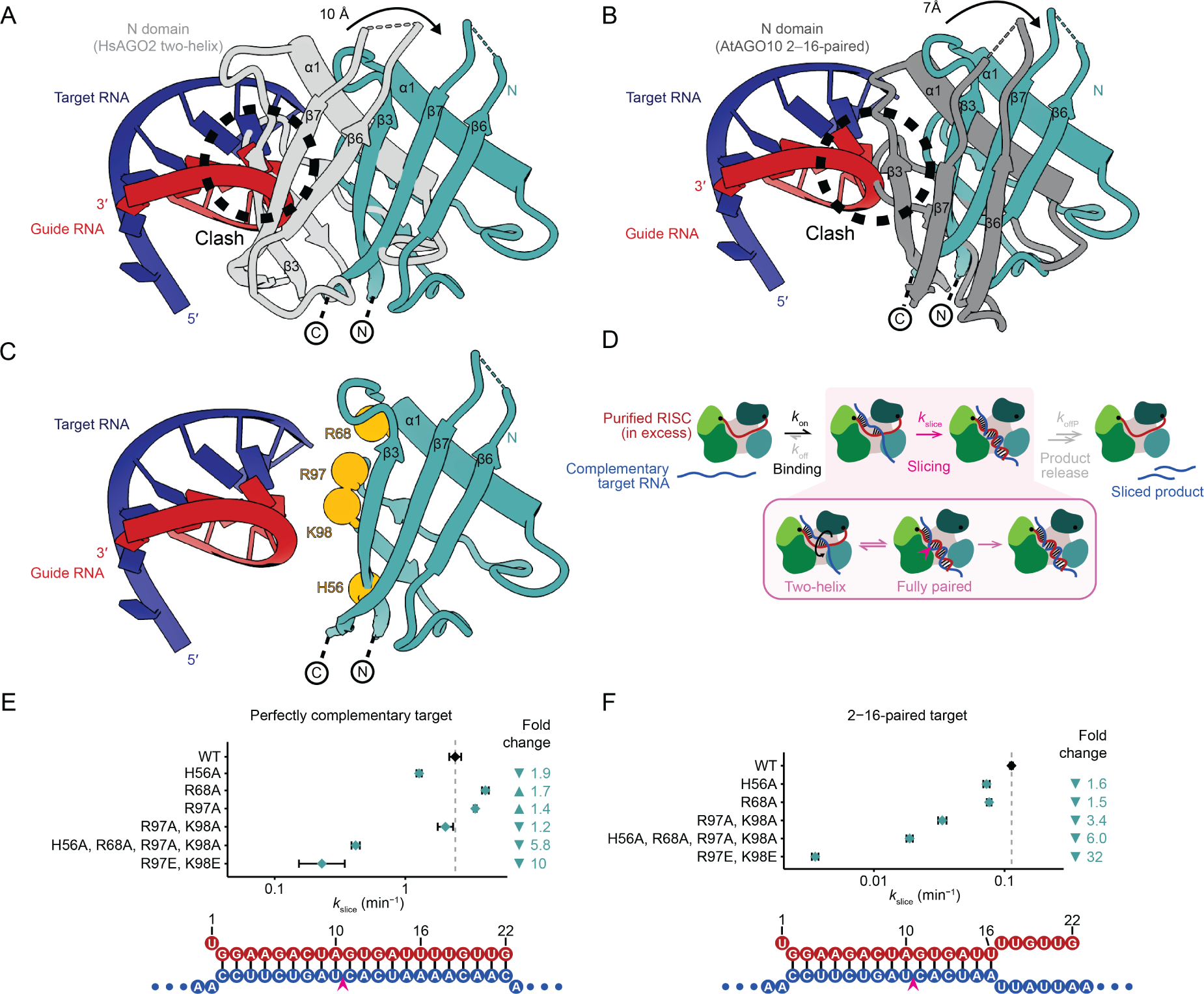
N-domain movements required to accommodate the fully paired guide−target RNA duplex. (A) Overlay of the N domain in the two-helix conformation (gray, PDB: 6N4O^31^) and the fully paired conformation (cyan, this study), front view. Guide and target nucleotides 15−22 are shown, with guide RNA colored red and target RNA colored blue. Models were aligned on the static MID and PIWI domains. The N domain moves by ∼10 Å between the two conformations to accommodate the fully paired RNA. (B) Overlay of the N domain in the AtAGO10 2−16-paired structure (dark gray, PDB: 7SWF^34^) and the fully paired conformation (cyan, this study), front view. Otherwise, this panel is as in **A**. The N domain moves by ∼7 Å between the two conformations to accommodate the fully paired RNA. (C) N-domain basic residues (show as orange spheres) that lie in close proximity to the RNA duplex, potentially forming contacts with backbone phosphates. Other colors are as in **A**. (D) Kinetic scheme of RISC-catalyzed slicing. The inset highlights the change from the two-helix conformation to the fully paired conformation, prior to slicing. For single-turnover reactions, elementary rate constants for target dissociation (*k*_off_) and product release (*k*_offP_) have negligible impact because RISC concentrations are in large excess over target, and slicing rates are instead a function of the elementary rate constants for target association (*k*_on_) and slicing (*k*_slice_). Otherwise, as in Figure 1A. (E−F) The *k*_slice_ values for N-domain mutants of HsAGO2–miR-7 RISC, slicing either a perfectly complementary target (**E**) or a target with mismatches after position 16 (**F**). Fold changes relative to wildtype (WT) protein (dashed line) are plotted above guide–target pairing schematics. Error bars indicate CIs from model fitting.

Superposing the extended RNA duplex of our HsAGO2 structure on the 2−16-paired AtAGO10 structure showed the expected clash with the N domain of AtAGO10 at position 17 of the duplex (**Figures 2B and S1D**). To accommodate a longer RNA duplex in the central channel, α1 moves by ∼7 Å between the AtAGO10 and HsAGO2 structures, and β3−β7, together with the remainder of the N domain, move by ∼7 Å (**Figure 2B**), allowing the RNA duplex to extend unencumbered alongside the N domain and then exit the central channel.

Our structure of HsAGO2 in the slicing-competent conformation, considered together with previous biochemical and comparative studies, suggested that contacts between the N domain and the RNA duplex might influence AGO-catalyzed slicing. From our structure, we identified residues H56, R68, R97, and K98 of the N domain as potentially contacting RNA backbone phosphates (**Figure 2C**). H56 is substituted to leucine in HsAGO3, a paralog with limited slicing activity (**Figure S4A**),^49^ and is located within the region (residues 28−64) shown in domain-swapping experiments to be essential for restoring full activity to HsAGO3.^42^ Importantly, three of these residues that orient toward the central channel (H56, R97, and K98) do not contact RNA in two-helix conformations that precede the slicing conformation,^31,48^ which suggests that interactions between these N-domain residues and RNA might favor slicing by stabilizing the slicing-competent conformation.

To test the impact of these residues on slicing, we purified miR-7–HsAGO2 complexes harboring point substitutions at these residues and examined their effects on slicing complementary targets. For each substitution, we focused on its impact on *k*_slice_, the elementary rate constant of slicing, reasoning that comparing *k*_slice_ values would capture any effects on either the chemical step of slicing (if it is rate limiting) or any rate-limiting, slicing-associated, conformational change that might be required after target binding and before the chemical step (**Figure 2D**). To obtain *k*_slice_ values, slicing kinetics were measured over a range of miR-7 RISC concentrations, but with RISC always in large excess over labeled target (**Figure S4B**). The range of RISC concentrations enabled *k*_slice_ to be disentangled from the elementary rate of target association (*k*_on_), and the excess RISC ensured single-turnover kinetics, which prevented potential confounding effects of any post-slicing steps, such as product release (**Figure 2D**).^32^

We first examined the impact of N-domain point substitutions on the slicing of a fully complementary target, for which the chemical step is rate limiting.^32^ In this context, single alanine substitutions of H56, R68, or R97, or a double substitution of R97 and K98 to alanines, each had minor impacts on slicing, either increasing or decreasing *k*_slice_ by no more than two-fold (**Figures 2E and S4B**). Nonetheless, substituting all four residues to alanine (H56A, R68A, R97A, K98A) reduced *k*_slice_ by 5.8-fold (95% confidence interval [CI], 5.1−6.6-fold), and more severe, charge-reversal substitutions of R97 and K98 to glutamates (R97E, K98E) reduced *k*_slice_ 10-fold (CI, 6.9−16-fold) (**Figures 2E and S4B**), consistent with the idea that contacts to the N domain enhance the chemical step of slicing. No substantial changes in binding kinetics (*k*_on_) were detected for these mutants, indicating that they were otherwise functionally intact (**Figure S4B**).

We next examined the impact of N-domain point substitutions on the slicing of a target mismatched at tail positions 17–22. For most guide RNAs, including miR-7, these mismatches to the tail reduce the occupancy of the centrally paired, slicing-competent conformation, such that occupancy of this centrally paired conformation joins the chemical step as rate-limiting for *k*_slice_.^32^ Single alanine substitution of H56 or R68, or a double substitution of R97 and K98 to alanines, was sufficient to reduce *k*_slice_ 1.6-fold (CI, 1.5−1.7-fold), 1.5-fold (CI, 1.4−1.6-fold), and 3.4-fold (CI, 3.1−3.7-fold), respectively (**Figures 2F and S4B**). The impact of the quadruple alanine substitution was retained (6.0-fold; CI, 5.7−6.4-fold), and the effect of charge-reversal substitutions at R97 and K98 was enhanced to 32-fold (CI, 30−34-fold) (**Figures 2F and S4B**). These results suggest that contacts between the N domain and the guide–target duplex enhance target slicing, especially the slicing of tail-mismatched targets, implying that RNA−protein contacts involving the N domain might work in concert with RNA base-pairing interactions to stabilize formation of the centrally paired, slicing-competent conformation. Contacts between a repositioned N domain and a fully paired guide– target duplex, although not homologous to those observed for HsAGO2 RISC, are also observed in recent structures of bacterial RISCs.^37,38^

Although flexibility of the N domain prevented modeling of these side chains contacting the distal region of the duplex (**Figures 2C, S3C, and S3E)**, our slicing analyses of point substitutions supported their function (**Figures 2E and 2F**). Moreover, these N-domain residues that influence slicing are largely conserved in animal and plant clade I−II AGOs (**Figure S4A**), suggesting that they have similar roles in other eukaryotic AGO proteins that catalyze slicing.

Taken together, our structural and mutagenesis results indicated that as the supplementary helix rotates into the central channel, the N domain also moves, achieving two aims. The first is to enlarge and vacate the central channel, creating space for the centrally paired duplex. The second is to form new contacts with the RNA duplex once it occupies the channel—contacts required for rapid slicing of the target transcript.

### The PAZ and linker domains rearrange to help form the slicing conformation

When transitioning from the two-helix conformation to the fully paired conformation, the PAZ domain must release the 3′ end of the guide (**Figure 1A**). Indeed, in our model of the slicing conformation, the 3′ end of the guide and its binding pocket within the PAZ domain have moved ∼63 Å from each other. This release untethers the 3′ end of the guide and the PAZ domain from each other. The PAZ domain of our structure exhibits increased flexibility and moves by ∼18 Å compared to the two-helix HsAGO2 structure^31^ (**Figure 3A**). This movement is ∼13 Å beyond the ∼5-Å movement observed in the AtAGO10 2−16-paired structure, in which the 3′ end of the guide is also released^34^ (**Figure 3B**). This further movement is presumably a consequence of additional structural remodeling required to accommodate a longer, fully paired duplex.

**Figure 3.**
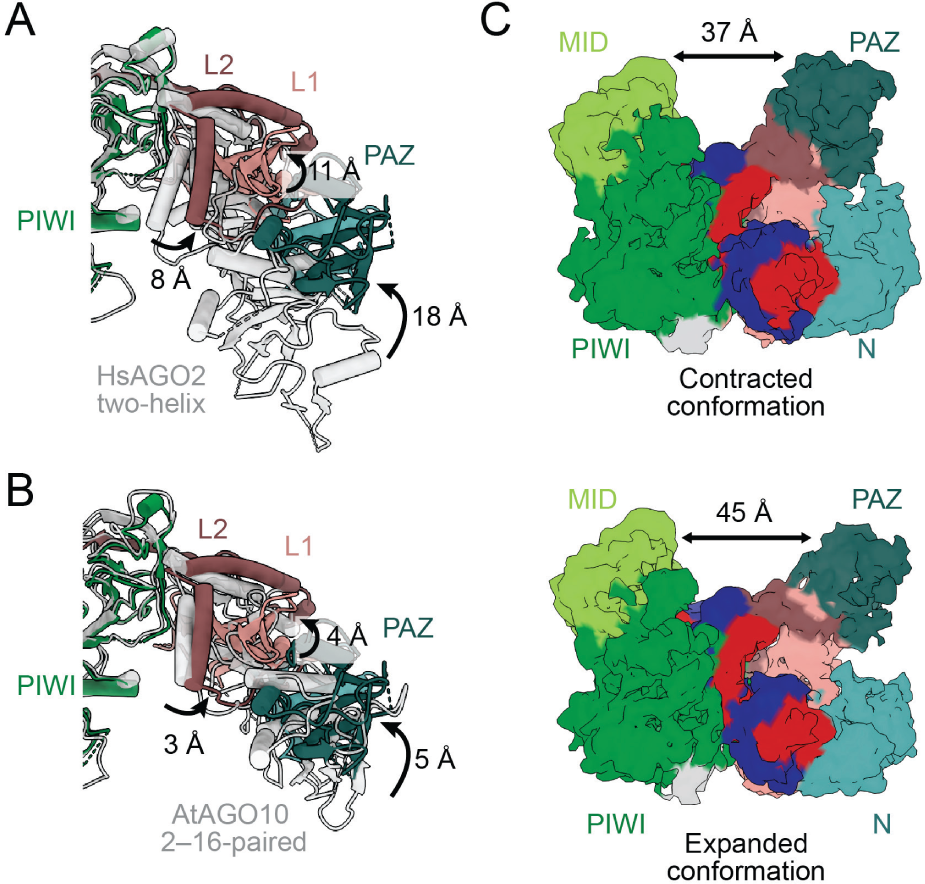
Flexibility of the N−PAZ lobe. (A) Movements of PAZ, L1, and L2 domains associated with the transition from the two-helix conformation (light gray, PDB: 6N4O^31^) to the fully paired conformation (colored as in Figure 1C, this study). A top view is shown. For clarity, the RNA duplex is not shown. (B) Movements of PAZ, L1, and L2 domains relative to the AtAGO10 2−16-paired conformation (light grey, PDB: 7SWF^34^); otherwise, as in **A**. (C) The central channel adopts relatively contracted and expanded conformations in the fully paired conformation. HsAGO2 and RNA are colored as in Figure 1D. The contracted and expanded conformations correspond to frames 0 (top) and 39 (bottom), respectively, of a 3DFlex movie. Models for all movie frames are provided (**Figure S5A**).

The L1 and L2 linker domains were rearranged in conjunction with the structural transitions of the N and PAZ domains. L1 acts as a hinge domain between the N−PAZ lobe and the MID−PIWI lobe.^16^ We observed rotation involving β10−β12 and α3 of L1, with movement of about 11 Å when compared to the two-helix conformation of HsAGO2^31^ (**Figure 3A**). This 11-Å movement is 7 Å beyond the 4-Å movement observed for α3 of L1 in the AtAGO10 2−16-paired structure (**Figure 3B**).

L2 connects the PIWI domain with the N domain and contains S387, a phosphorylation site reported to regulate slicing.^50^ Residues 406−444 of L2, which interact with the PIWI domain (and position 1 of the target), remain fixed; the other half of L2, which includes helices α7−α9, moves by ∼8 Å relative to the two-helix conformation^31^ (**Figure 3A**). This 8-Å movement is 5 Å beyond the 3-Å movement observed in the L2 domain in the AtAGO10 2−16-paired structure (**Figure 3B**).

The conformational changes observed in L1 and L2 appear to accommodate the movement of the supplementary helix as it rotates into the widened central channel.

More broadly, when considered in the context of the series of changes that occur over the four steps of target pairing, these changes at step four continue the monotonically increasing separation between the MID−PIWI and N−PAZ lobes, with widening of the central channel. This widening successively increases the distance between residues of the central gate (residues 353−358 and 602−608 of L2 and PIWI, respectively^31^), from 8 Å without bound target to 9, 14, and 25 Å after steps two, three, and four, respectively (**Figure 1A**).^19,30,48^ Thus, these observations extend previous analyses indicating that binding of target RNAs with increasing complementarity entails successive opening of the protein, achieved by movement of the N, L1, PAZ, and L2 domains.^19,31,48^

### HsAGO2 is conformationally dynamic through mobility of the N and PAZ domains

To further characterize the implied flexibility in the N and PAZ domains, we performed 3DFlex analysis as implemented in cryoSPARC^51^ (**Figure S2C and S5A**). This analysis indicated that the MID and PIWI domains remain relatively static, whereas the N and PAZ domains appear to move across a range of states (**Movie S3**). This motion explains the lower occupancy observed for the N and PAZ domains in our maps. The first and last frames of the 3DFlex movie revealed contracted and expanded conformations of the complex, in which the N and PAZ domains are positioned closer to or further from the RNA duplex, respectively (**Figures 3C and S5A**). Between the two conformations, the N domain moves by ∼4 Å and the PAZ domain moves by ∼8 Å (**Figure 3C**). In conjunction, the RNA appeared the most mobile at positions 5–12 of both strands (**Movie S3**).

### HsAGO2 makes extensive contacts with the RNA duplex in the slicing conformation

Analysis of our model indicated that, when in the slicing-competent conformation, HsAGO2 makes many contacts to the RNA duplex (**Figures 4A−D**). Most of these contacts are analogous to those observed for AtAGO10 in the 2−16-paired conformation.^34^ With respect to a key interaction between R179 (R288 in AtAGO10), which resides in an L1 β-hairpin, and centrally paired target RNA,^34^ the backbone density of our map supports this proposed contact, although side-chain density was not sufficiently resolved for it to be directly observed (**Figures 4A−C**). In addition to these orthologous contacts, we also propose interactions between the remodeled N domain, which is unique to our structure, and positions 15–19 of the duplex, some of which are also unique to our structure (**Figures 4A−C**).

**Figure 4.**
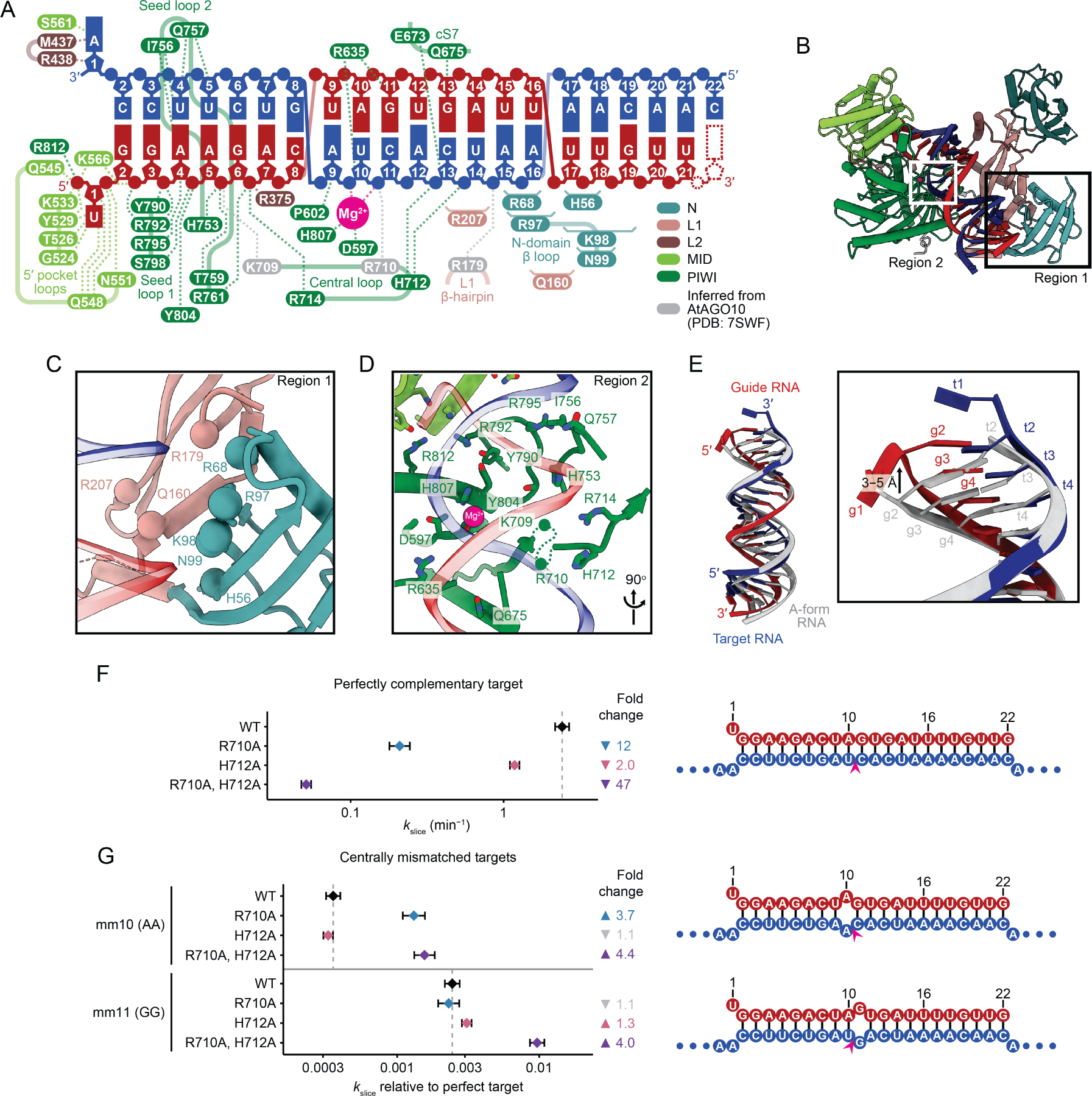
HsAGO2−RNA contacts involved in slicing. (A) Schematic of HsAGO2–RNA contacts in the fully paired, slicing-competent conformation. Colors are as in Figure 1B **and 1C**. Colored dotted lines represent protein–RNA interactions observed in our structure. Brackets represent inferred interactions involving poorly resolved side chains. Gray dotted lines represent interactions analogous to those observed in the AtAGO10 2−16-paired structure (PDB: 7SWF^34^) and, although consistent with our structure, not observed in our structure due to poorly resolved side chains. Guide position 22 (dotted) is poorly resolved but implicated as paired to target position 22. (B) Model of the fully paired conformation of HsAGO2 RISC with region 1 (containing the N and L1 domains) and region 2 (containing the active site and central-loop) highlighted by boxes. Colors are as in Figure 1E. (C) Protein–RNA contacts of the N and L1 domains observed in the fully paired conformation of HsAGO2 RISC (region 1). A front view is shown, with colors as in Figure 1E. For clarity, RNA nucleobases are not shown. Interacting residues are shown as spheres due to insufficient side-chain density. (D) Protein–RNA contacts of the PIWI domain, particularly around the seed region, active site, and central loop, observed in the fully paired conformation of HsAGO2 RISC (region 2). A rotated front view is shown, with colors as in Figure 1E. For clarity, RNA nucleobases are not shown. Interacting residues lacking density are shown as spheres along dotted lines. (E) Distortions from an A-form helix observed for the guide−target RNA duplex (**Movie S4**), culminating in a 3–5 Å displacement of guide position 2 (g2, arrow in inset). A standard A-form duplex (gray) is superimposed after alignment at positions 7−12 of the guide strand. Colors are as in Figure 1B. (F) *k*_slice_ values for wildtype (WT) or central-loop mutants of HsAGO2–miR-7 RISC paired with the perfectly complementary slicing target (pairing schematic shown on the right). Otherwise, this panel is as in Figure 2E. Wildtype value is replotted from Figure 2E. (G) *k*_slice_ values for wildtype (WT) or central-loop mutants of HsAGO2–miR-7 RISC paired with targets harboring a central mismatch at either position 10 (mm10) or 11 (mm11). Values are reported relative to those of perfectly complementary targets. Fold changes that were not statistically significant are shown in gray. Otherwise, this panel is as in **F**.

In the seed region, most of the contacts are to the guide strand (**Figure 4A**). Many involve basic side chains contacting guide backbone phosphates, but other types of interactions are also observed, including hydrogen bonds to riboses, and hydrogen bonds and hydrophobic interactions with nucleobases at target positions 4−5, in the minor groove. These contacts to the seed region are largely consistent with those reported for the seed-paired conformation of HsAGO2 RISC,^19,31^ which supports the idea that the seed region remains stationary as the rest of the RNA and protein are structurally remodeled to reach the fully paired conformation.

Presumably because of these many interactions with the seed region, this region of the RNA duplex deviates the most from standard A-form helix. Compared to A-form, positions 2−4 of the guide extend along the helical axis by ∼3−5 Å without commensurate movement of the target, thereby lengthening the helix by the equivalent of an additional base pair, with the differential movement of the two strands presumably flattening the base-pair inclination angles in this region^52^ (**Figure 4E**). Examination of models of other eukaryotic RISC–target structures revealed a similar phenomenon.^31,34,48^ Other than a slight widening of the major groove in the central region and a narrowing more distally, the remainder of the duplex resembled standard A-form geometry (**Figure S4C, Movie S4**). This is in contrast with the model of 2−16-paired AtAGO10,^34^ which has more pronounced widening of the major groove in the central region (**Figure S4C, Movie S4**).

In the central region, contacts to the RNA are primarily to the target strand (**Figures 4A, 4B, and 4D**). We observed two catalytic residues (D597 and H807) coordinating a Mg^2+^ ion that was also coordinated to the scissile phosphate in the active site, analogous to the plant and bacterial structures.^33,34^ The second active-site Mg^2+^ ion was not observed, as expected because D669, which typically coordinates the second active site Mg^2+^ ion,^33^ was mutated to create a catalytically dead enzyme (**Figure 4D**). The fourth catalytic residue, E637, is in the active, plugged-in position as expected for HsAGO2 and for the slicing-competent conformation (**Figure S4E**).^26^

A conserved, highly basic loop spanning HsAGO2 residues 709–714, which we term the central loop, lies in the middle of the central channel, adjacent to the active site (**Figure 4D**). This loop, together with the L1 β-hairpin and the cS7 (conserved-sequence 7) loop in the PIWI domain,^53^ forms the active-site channel previously described in the centrally paired AtAGO10 structure.^34^ Side chains of central-loop residues K709 and R710 were omitted from our model due to lack of density, although both are resolved in the AtAGO10 structure.^34^ Previous analyses indicate that K709 and R714 interact with backbone phosphates of the guide at seed positions 6 and 7, respectively—interactions observed even before initial binding of target (**Figure S4D**).^24,25^ In our model, R714 mediates the same interaction with position 7 of the guide (**Figures 4A, 4D, and S4E**). In contrast to these contacts to the seed region of the guide, the other basic residues of the central loop reach to phosphates in the central region of the target. Specifically, we observed that H712 interacts with the backbone phosphates of target positions 12 and 13 (**Figures 4A, 4D, and S4E**), and analogy to the AtAGO10 structure suggests that R710 (R836 in AtAGO10) interacts with the scissile phosphate at target position 10 (**Figures 4A and S4D**).^34^ These latter interactions involving R710 and H712 can form only upon completion of central pairing and are thus unique to the slicing-competent conformation. As with all other residues of the central loop, R710 and H712 are essentially invariant in sequenced animal AGOs and well conserved in plants (**Figure S4A**). Together, RNA interactions of the central loop contact and straddle the scissile phosphate, spanning the major groove to contact backbone phosphates of both the seed of the guide RNA and the central region of the target RNA, on opposite sides of the active site. We propose that these contacts help position centrally paired RNA in the active site for slicing.

To investigate the impact of central-loop residues, we examined slicing activity of miR-7 RISC with alanine substitutions in this loop. Substituting H712 with alanine reduced *k*_slice_ by 2.0-fold (CI, 1.8−2.3-fold), whereas substituting R710 reduced *k*_slice_ by 12-fold (CI, 9.6−14-fold) (**Figures 4F and S4F**). Substituting both R710 and H712 reduced *k*_slice_ 47-fold (CI, 41−54-fold) (**Figures 4F and S4F**), indicating that R710 and H712 both help establish slicing-compatible interactions around the active site in a non-redundant manner. Importantly, no substantial changes in binding kinetics (*k*_on_) were detected for these mutants (**Figure S4F**), which supports the model that these interactions mediated by R710 and H712 are unique to the slicing-competent conformation and occur after nucleation of binding at the seed region (steps one and two, **Figure 1A**).

R710 and H712 are poorly conserved in the central loops of PIWI proteins and prokaryotic AGOs (**Figure S4A**), which have less stringent central base-pairing requirements for slicing.^54,55^ We hypothesized that the loss of these contacts might explain why slicing by these other proteins is more permissive. To test this idea, we examined the reduction in *k*_slice_ upon introduction of either an A:A RNA mismatch at position 10 or a G:G mismatch at position 11, in the context of different central-loop mutants. Substituting both R710 and H712 with alanines reduced the loss of *k*_slice_ caused by either RNA mismatch, by 4.4-fold (CI, 3.6−5.4-fold) and 4.0-fold (CI, 3.4−4.7-fold) respectively (**Figures 4G and S4G**). The impact of single substitutions was dependent on the specific RNA mismatch. Substituting R710 (which interacts with target position 10) with an alanine reduced the loss of *k*_slice_ due to a position-10 mismatch by 3.7-fold (CI, 3.0−4.6-fold), whereas substituting H712 (which interacts with target positions 12−13) with an alanine reduced the loss of *k*_slice_ due to a position-11 mismatch by 1.3-fold (CI, 1.1−1.5-fold) (**Figures 4G and S4G**). These results indicate that the central loop might partially explain the greater stringency for pairing near the active site observed for metazoan AGOs.

### A PIWI loop distant from the target RNA regulates its dissociation

When modeling the PIWI domain, we observed additional density that appeared to represent the eukaryotic insertion (EI) loop at residues 820−837. The structure of this loop is not modeled in any previous structural study of HsAGO2, presumably because of its flexibility. The EI loop is highly conserved among metazoan AGOs and is reported to regulate target release through the phosphorylation of five closely spaced phosphorylation sites by casein kinase 1α (CK1α),^56,57^ with the proposal that direct electrostatic repulsion between negative charges of the phosphorylated EI loop and the phosphate backbone of target RNA promotes target release.^58^

Using the additional density in our data and an AlphaFold2 model^59^ of HsAGO2, we modeled the full EI loop to ∼7 Å local resolution (**Figure S5B**). Superimposing our model of the EI loop with a model of HsAGO2 bound to target with supplementary pairing^31^ indicated that the phosphorylated residues are positioned too far away (>15 Å) from the target RNA backbone to readily mediate direct electrostatic repulsion (**Figure S5B**). Nonetheless, our biochemical analyses confirmed that phosphomimetic substitutions to glutamates at all five phosphosites in the EI loop caused faster release of targets with supplementary pairing (**Figures S5C−E**), in agreement with previous results using in vitro-phosphorylated HsAGO2.^58^ Indeed, when associated with a more optimal guide and target, HsAGO2 with phosphomimetic substitutions in the EI loop released target 20-fold (CI, 17−25-fold) faster than wildtype HsAGO2 (**Figures S5C−E**), bringing to the fore the question of the structural basis for this activity.

## DISCUSSION

Our structure provides a missing piece of the AGO catalytic cycle, showing how a eukaryotic AGO accommodates a fully base-paired guide−target duplex to support slicing. While representing a later step of the target-slicing pathway, it likely also represents an early step in AGO-loading pathway—specifically, the slicing-dependent loading that occurs for siRNA duplexes and some miRNA duplexes that have passenger strands with perfect (or near-perfect) complementarity.

As the complex proceeds through the four steps of target pairing (**Figure 1A**), the major movements of the N, L1, PAZ, and L2 domains each build on the previous movements (**Movie S2**)—with successive movements each further widening the RNA-binding channel, as expected if each conformation facilitates the subsequent pairing needed to progress to the next conformation. This phenomenon continues a theme first observed for HsAGO2 RISC when studying pairing to the full seed, which induces changes proposed to facilitate subsequent pairing to the supplementary region.^19^

Our results also speak to the stability of the slicing-competent conformation of RISC. Previous crystallographic results^31^ have been interpreted to suggest that only a small fraction of substrate-bound HsAGO2 RISC populates the fully paired, slicing-competent conformation. However, in our cryo-EM analysis, we observed only the slicing-competent conformation. This observation supports the idea that, upon binding to fully paired targets, HsAGO2 RISC, and presumably orthologs in other species, predominantly populates the slicing-competent conformation. This idea also concurs with results of chemical footprinting and single-molecule FRET experiments.^32,36^

We suggest two explanations for why we observed only the slicing-competent conformation, whereas previous structural studies observed other conformations. (1) Previous attempts to determine the structure of eukaryotic RISC in a centrally paired, slicing-competent conformation, which include a crystallographic analysis of HsAGO2^31^ and a cryo-EM analysis of AtAGO10^34^, used target RNAs that pair up to position 16 and no further. The crystallographic study of HsAGO2 yielded only a two-helix conformation, whereas the cryo-EM analysis of the AtAGO10 ternary complex yielded a mixture of a two-helix and a single-helix conformation. We did not observe any two-helix conformation in our analysis of RISC with a fully complementary target. This outcome, combined with recent biochemical insight, suggests that the previous structural efforts capture these stalled states due to the use of target RNAs that do not pair beyond position 16, in that HsAGO2 RISC bound to such targets does not as readily populate a centrally paired conformation.^32^ (2) Inspection of crystal-packing interactions in previous HsAGO2 X-ray crystal structures suggests that these interactions would constrain the mobility of the N and PAZ domains, preventing movements of these domains required to accommodate a fully paired duplex (**Figure S1A**). Crystal packing interactions appear to also limit mobility of the N and PAZ domains of TtAGO^35^ (**Figure S1B**). Moreover, if the two-helix conformation is more static than the centrally paired conformation, crystallization might preferentially capture it, even if it is not the most populated conformation. In contrast, cryo-EM can capture the more flexible slicing-competent conformation.

We obtained evidence that the slicing-competent conformation moves between contracted and expanded states (**Movie S3**). Likewise, smFRET studies indicate that when HsAGO2 is bound to a fully complementary target, it takes on more dynamic conformations in solution.^36^ The movements we observe might subtly alter the coordination geometry of the active site and might be sensitive to nucleotide sequence, in which case, they could help explain sequence features associated with more rapid slicing.^32^ The contracted and expanded states might also represent HsAGO2 poised for different stages of the slicing process, such as endonucleolytic cleavage and product release, respectively.

## ACKNOWLEDGMENTS

We thank K. Heindl, M. Herzik, L.K. Kinman, D.H. Lin, M. Wilkson, and members of the Bartel and Vos laboratories for helpful discussions. Cryo-EM specimens were prepared and data was collected at the Cryo-EM facility at MIT.nano. We thank C. Borsa, S. Sterling and J. Podgorski for support at MIT.nano. A.A.M. is supported by the National Science Foundation Graduate Research Fellowship Program. P.Y.W. is supported by the MIT Office of Graduate Education Fellowship. This work was supported by NIH grant GM118135 (to D.P.B.). D.P.B. is an investigator of the Howard Hughes Medical Institute. S.M.V. is a Freeman Hrabowski Scholar of the Howard Hughes Medical Institute. Research in the Vos Lab is supported by the Smith Family Foundation, Alex’s Lemonade Foundation Crazy Eight Initiative, and the NIH Director’s New Innovator Award (DP2-GM146254).

## AUTHOR CONTRIBUTIONS

A.A.M. prepared cryo-EM grids, collected and processed the cryo-EM data, and built and analyzed the structure model. P.Y.W. prepared samples of the HsAGO2 ternary complex for cryo-EM, conducted biochemical experiments and associated data analyses, and assisted with model analysis. D.P.B. and S.M.V. supervised the study. All authors wrote the manuscript.

## STAR METHODS

### Data availability statement

Coordinates are deposited in the PDB (9CMP), and maps are deposited in the EMDB (EMD-45752, EMD-45753). All code used for analysis and graphing of biochemical data is available at https://github.com/wyppeter/AGO2_slicing2_2024. Plasmids generated in this study will be deposited at Addgene. Gel images, reaction assay data, and multiple sequence alignments will be deposited at a publicly available repository.

### Method details

#### Purification of SENP^EuB^ protease

SENP^EuB^ protease was purified as described.^60^ Briefly, Rosetta 2(DE3) *E. coli* cells (MilliporeSigma, 714003) were transformed with the pAV0286 plasmid (Addgene, #149333), plated on LB agar plates with kanamycin and chloramphenicol, then grown in Terrific Broth with kanamycin, chloramphenicol, and Antifoam SE-15 (Sigma Aldrich, A8582) at 37°C to reach 0.1−0.2 OD, then at 18°C to 0.6−0.8 OD, followed by overnight induction of expression with 0.2 mM IPTG. Cell pellets were collected by centrifugation at 4000*g* at 4°C for 15 min, resuspended in lysis buffer (50 mM Tris pH 7.5, 300 mM NaCl, 5 mM imidazole, 4 mM β-mercaptoethanol, 1 tablet per 10 mL cOmplete Mini EDTA-free Protease Inhibitor Cocktail [Roche, 11836170001]), and lysed by sonication. Cell lysate was clarified by centrifugation at 40000*g* at 4°C for 30 min. SENP^EuB^ was purified from the clarified lysate using Ni-NTA Agarose (Qiagen, 30210) and eluted with lysis buffer with 250 mM imidazole, without protease inhibitors. The eluate was dialyzed overnight at 4°C into gel filtration buffer (50 mM Tris pH 7.5, 150 mM NaCl, 4 mM β-mercaptoethanol), then purified by gel filtration on a Superdex 200 Increase 10/300 GL column (Cytiva, 28990944). Peak fractions were collected, and 80% (v./v.) glycerol was added to bring glycerol up to 10% (v./v.) final concentration. Aliquots were flash frozen in liquid nitrogen for storage at −80°C.

#### Preparation of biotinylated target RNA

The target RNA had full complementarity to the 22-nt isoform of miR-7. This complementary site was flanked by three 2′-*O*-methylated uridines on its 5′ terminus and two 2′-*O*-methylated adenines on its 3′ terminus, followed by a deoxyuridine with a 5-octadiynyl modification at the 3′ end (**Table S1**). This oligonucleotide was synthesized (IDT), gel purified on a 15% urea-polyacrylamide gel, and resuspended in water. Biotin was added to the 3′ end by click chemistry, incubating 100 µM gel-purified oligonucleotide with 2 mM diazo biotin azide (Click Chemistry Tools, 1041-25), 50 mM sodium phosphate pH 7.0, 5 mM sodium ascorbate, 2.5 mM THPTA (Click Chemistry Tools, 1010-100), and 2.5 mM copper (II) sulfate, at 30°C for 20 min. The reaction was mixed with 0.6X volume of gel-loading buffer (8 M urea, 25 mM EDTA, 0.025% [w./v.] xylene cyan, 0.025% [w./v.] bromophenol blue), then purified on a 15% urea-polyacrylamide gel and resuspended in water.

#### Purification of the HsAGO2^D669A^−miR-7−target ternary complex for cryo-EM

The pcDNA3.3−3xFLAG−SUMO^Eu1^−HsAGO2^D669A^ plasmid was cloned from pcDNA3.3−3xFLAG−HsAGO2^D669A^ (Addgene plasmid #220461) and pAV0279 (Addgene plasmid #149688)^60^ by restriction cloning, using NheI and BamHI sites introduced by PCR with KAPA HiFi HotStart Ready Mix (Roche, 07958935001) (**Table S1**). Sequences encoding GGAS and GS linkers were used to join 3xFLAG and SUMO^Eu1^, and SUMO^Eu1^ and HsAGO2^D669A^ sequences, respectively. Plasmids were verified by Sanger sequencing and prepared using the PureLink HiPure Expi Plasmid Gigaprep Kit (Invitrogen, K210009XP).

5′-phosphorylated, 22-mer miR-7 guide and passenger RNAs (**Table S1**) were synthesized (IDT), purified on a 15% urea-polyacrylamide gel, and then resuspended in water. The guide−passenger RNA duplex was annealed using 25 μM of each RNA in annealing buffer (60 mM Tris-HCl pH 7.5, 200 mM NaCl, and 2 mM EDTA), which was incubated at 90°C and then slowly cooled over 1.5 h to 30°C, before being chilled on ice and stored at −80°C.

2 L of Expi293F cells (Gibco, A14527), cultured in Expi293 expression media (Gibco, A1435102) at 37°C in 5% CO_2_, were transfected at a density of ∼2 million cells per mL, with 1.9 mg of pcDNA3.3−3xFLAG−SUMO^Eu1^−HsAGO2^D669A^, 0.1 mg of pMaxGFP (Lonza), and 6 mg of polyethylenimine (Polysciences, 23966) incubated in 100 mL of Opti-MEM (Gibco, 51985091) at room temperature for 20 min. After culture for 40 h at 37°C in 5% CO_2_, cells were harvested and lysed in 80 mL of hypotonic buffer (10 mM Tris pH 7.5, 10 mM KOAc, 1.5 mM Mg(OAc)_2_, 2% [v./v.] glycerol, 0.5 mM TCEP, 5 mM NDSB-256 [MilliporeSigma, 48-001-05GM], with cOmplete EDTA-free Protease Inhibitor Cocktail [1 tablet per 50 mL, Roche, 11873580001]) by a Dounce homogenizer. Cell lysate was clarified by centrifugation at 3000*g* at 4°C for 15 min, then again for 30 min. The supernatant was re-equilibrated by adding 25% volume of re-equilibration buffer (150 mM Tris pH 7.5, 700 mM KOAc, 15 mM Mg(OAc)_2_, 42% [v./v.] glycerol, 0.5 mM TCEP, 5 mM NDSB-256, with 1 tablet per 10 mL cOmplete Mini EDTA-free Protease Inhibitor Cocktail), before further clarification by centrifugation at 60,000*g* at 10°C for 20 min.

The resultant cell lysate was incubated with 50 nmol of annealed miR-7 guide−passenger RNA duplex for 1 h at 25°C on a gentle rocker. The incubated lysate was re-clarified by centrifugation at 6000*g* at 4°C for 10 min, followed by filtration through 5 µm PVDF filters (Millipore, SLSV025LS). The clarified lysate was incubated with 10 mL of 50% slurry of anti-FLAG M2 agarose (Millipore, A2220), pre-equilibrated in equilibration buffer (50 mM Tris pH 7.5, 150 mM NaCl, 1 mM MgCl_2_, 10% [v./v.] glycerol, 0.5 mM TCEP) according to manufacturer’s instructions, at 4°C for 1 h with end-over-end rotation. The agarose was collected by centrifugation at 100*g* at 4°C for 3 min, resuspended with 30 mL of equilibration buffer, and transferred into a glass column (Bio-Rad, 7372512). The resin was washed at 4°C with 20 mL of equilibration buffer, 20 mL of high-ATP/Mg^2+^ buffer (30 mM Tris pH 7.5, 150 mM NaCl, 20 mM MgCl_2_, 10% [v./v.] glycerol, 5 mM ATP, 0.5 mM TCEP), 20 mL of EDTA/EGTA buffer (30 mM Tris pH 7.5, 150 mM NaCl, 10% [v./v.] glycerol, 1 mM EDTA, 1 mM EGTA, 0.5 mM TCEP), and 30 mL of equilibration buffer. To bind the target RNA, the resin was resuspended in-column with 3 nmol of biotinylated target RNA pre-warmed at 37°C for 2 min in 5 mL of equilibration buffer, and incubated at 25°C for 40 min. After flowthrough was drained from the re-settled resin, the resin was first washed by resuspending in-column in 20 mL of equilibration buffer at 25°C and incubating for 20 min. The resettled resin was washed again with 20 mL of equilibration buffer at 4°C. To elute the ternary complex, the resin was resuspended in 5 mL of equilibration buffer with 400 nM SENP^EuB^, and incubated at 4°C for 30 min. After collecting the first eluate, the resin was washed with 5 mL of equilibration buffer, and the wash was combined with the first eluate. The combined eluate was centrifuged at 6000*g* at 4°C for 10 min, and the supernatant was incubated with 300 µL of MagStrep Strep-Tactin XT beads (IBA Lifesciences, 2-4090-002) slurry pre-equilibrated in equilibration buffer, at 4°C for 1 h with end-over-end rotation. The beads were washed twice with 1 mL of equilibration buffer, then once with 1 mL of gel filtration buffer (50 mM Tris pH 7.5, 150 mM NaCl, 1 mM MgCl_2_, 2% [v./v.] glycerol, 0.5 mM TCEP). To elute the bound complex, the beads were incubated with 150 µL of biotin elution buffer (50 mM Tris pH 7.5, 150 mM NaCl, 1 mM MgCl_2_, 2% [v./v.] glycerol, 0.5 mM TCEP, 50 mM biotin), at 4°C for 1 h with gentle rotation. The biotin eluate was centrifuged at 17000*g* at 4°C for 10 min, and the supernatant, estimated at 1.3 µM based on absorbance at 280 nm, was used for gel filtration. 100 µL of complex at 1.3 µM was applied to a Superdex 200 Increase 3.2/300 column (Cytiva 28990946) equilibrated in gel filtration buffer on an AKTA Micro purification system (Cytiva) at 4°C. Peak fractions were analyzed by SDS-PAGE followed by Coomassie staining with Imperial Protein Stain (Thermo Scientific, 24615). Peak fractions corresponding to ternary complex were chosen for grid preparation.

#### Cryo-EM grid preparation

Grid preparation was optimized to yield uniform, monodisperse particles. We observed sample denaturation when freezing the complex in pure ice and with the addition of the detergent CHAPSO. To prevent sample denaturation, we employed support layers. Aggregation was observed on monolayer graphene grids, but monodisperse particles were observed on Cu300 R2/2 grids coated with a 2-nm layer of amorphous carbon (Quantifoil: Q3100CR2-2NM). The grids were glow discharged on a K100X glow discharger for 12 s at 25 mA and mounted in a Vitrobot Mark IV (FEI Company) maintained at 4°C and 100% humidity. 4 µL of complex was applied to the amorphous carbon layer and incubated for 30 s, after which the grid was manually blotted from the back side for 4 s and plunged into liquid ethane.

#### Cryo-EM collection and data processing

Cryo-EM data were collected on a FEI Titan Krios II transmission electron microscope operated at 300 keV. A K3 summit direct detector (Gatan) with a GIF quantum energy filter (Gatan) was operated with a slit width of 20 eV. Automated data acquisition was performed with FEI EPU v2.12.1 software at nominal magnification of 130,000x, corresponding to a pixel size of 0.654 Å per pixel. Data acquisition was done at a defocus range of 0.2–1.4 μm. Image stacks of 50 frames were collected over 1 s in counting mode. The total dose for the sample was 51.0 e^-^/ Å^2^. A total of 12,106 image stacks were collected.

Frames were stacked, motion-corrected, and the contrast transfer function was estimated. Particles were auto-picked and extracted using a box size of 320 pixels in Warp 1.1.0 beta^61^, yielding 1,427,326 particles. The particle stack was imported into cryoSPARC v4.2.0^62^. Particles were re-extracted at 300 pixels and an initial reference-free 2D classification was done with the following parameters (all other parameters were left at their default values): number of classes = 150, maximum resolution = 10, initial classification uncertainty factor = 1, circular mark diameter = 120, re-center mask threshold = 0.77, Force max over poses/shifts = off, number of online-EM iterations = 50, batchsize per class = 400. On the original particle stack five rounds of succussive reference-free 2D classification were performed with the following parameters (all other parameters were left at their default values): number of classes = 100, initial classification uncertainty factor = 1, circular mark diameter = 120, circular mask diameter outer = 140, Force max over poses/shifts = off, number of online-EM iterations = 50, batchsize per class = 5000. After every round of 2D classifications, classes were excluded in a conservative manner, only removing obviously non-aligning particles. After the fifth round of 2D classification, 808,413 particles remained. This particle stack was re-extracted and re-centered and then subjected to two more rounds of reference-free 2D classification with the same parameters and only well aligning particles were retained, resulting in a final particle stack containing 222,377 particles.

The resulting particle stack was used for ab-initio reconstruction^62^ as implemented in cryoSPARC. The following parameters were used (all other parameters were left at their default values): number of ab-initio classes = 6, maximum resolution = 6, initial resolution = 9, initial minibatch size = 300, final minibatch size = 1000. One of the six ab-initio classes (class #4) resembled the HsAGO2 ternary complex, corresponding to a particle stack of 58,375 particles. A Non-Uniform refinement^63^ job was performed with this particle stack and the ab-initio reconstruction volume as an input low-pass filtered to 12 Å and minimize over per-particle scale turned on. In addition, a Non-Uniform refinement job was performed with an input particle stack from class 4 and each other ab-initio class with the ab-initio reconstruction volume from class 4 as an input low-pass filtered to 12 Å and minimize over per-particle scale turned on. Combination and Non-Uniform refinement of class 4 with class 2 and class 4 and 3 resulted in two maps (with 89,722 and 91,651 particles respectively) of the HsAGO2 ternary complex lacking density for the PAZ domain and having weak density for the N domain.

To further resolve the N and PAZ domains in both maps, a focus mask around the N and PAZ domains was generated in ChimeraX^64^ and then imported into cryoSPARC where the map was low-pass filtered to 20 Å and a mask was created using a dilation radius of 5 Å and soft padding width of 3 Å. The solvent mask used in the 3D classification was generated around the entire map using the same parameters as the focus mask. Each Non-Uniform refined map (class 2 + 4 and class 3 + 4) were treated separately and 3D classified using the focus mask separately. 3D classification was done with the following parameters (all other parameters were left at their default values): number of classes = 2, target resolution = 5, number of O-EM epochs = 5, O-EM batch size (per class) = 5000, initialization mode = PCA, force hard classification = on, class similarity = 0.1. 3D classification resulting in two classes, one with defined density for the N and PAZ domains, corresponding to a particle stack of 44,267 for the first map and 43,664 for the second map. These particles were subjected to Non-Uniform refinement (low-pass filtered to 12 Å and minimize over per-particle scale applied). This resulted in the highest resolution for the N and PAZ domains for each map, originating from the maps generated from ab-initio class 2 + 4 and 3 + 4. At this point, duplicate particles were removed and homogenous reconstruction was performed on the particle stack to merge the two particle sets. A final non-uniform refinement was done (low-pass filtered to 12 Å and minimize over per-particle scale applied) which resulted in a final particle stack of 69,008 particles. This resulting map was sharpened with a B-factor of –100 Å, resulting in Map 1, and local resolution estimation was determined using the built-in local resolution estimation tool in cryoSPARC. To mitigate orientation bias, we employed the anisotropy correction module of spIsoNet.^47^ The anisotropy correction module of spIsoNet was run with default parameters. The resulting half maps were post processed in Relion5^65^ and sharpened with a B-factor of –100 Å, resulting in Map 2.

3DFlex^51^ analysis was employed in cryoSPARC v4.4.0. Using the particle stack and volume from Map 1 a 3DFlex Data Preparation job was run with default parameters, followed by a Flex Mesh Preparation job with default parameters except for mask threshold set to 0.4 and stiffen low density regions turned on. A 3DFlex Training job was run with default parameters, and the 3DFlex Generator job was used to generate the volume series, with 41 frames per series. A movie of the frames was generated in Chimera.^66^

#### Model building

An X-ray crystal structure of HsAGO2 (PDB: 4OLA^24^) was rigid body fit into Map 2. The model was split into three different pieces consisting of residues 23−174, residues 175−412, and residues 413−856, and each piece was individually rigid body fit using ChimeraX.^64^ To model the RNA, positions 1−8 for the guide and target were used from a previously determined X-ray crystal structure (PDB: 6N4O^31^). To model positions 9−22 of the RNA, an A-form RNA duplex was generated in ChimeraX. Both pieces of the RNA were rigid body fit in ChimeraX. The model was manually adjusted in Coot v0.9.8.91.^67^ The RNA duplex was manually adjusted in Coot by setting weight restraints at 5.0 and performing an all-atom refinement. Further manual adjustment of the RNA model was done to satisfy base pairing and the placement of the sugar−phosphate backbones into the density. To better fit the PAZ domain into the density, a 30 second simulation was done in ISOLDE^68^ and then imported back into Coot and manually adjusted.

The model was real-space refined in Phenix version 1.20.1-4487-000.^69^ Ready Set was used to prepare input PDB models for refinement. Real-space refinement was performed with one macro-cycle of global minimization and ADP refinement, with a nonbonded weight of 2000 and an overall weight of 0.5. Information about the regions modeled are shown in **Table S2**. Statistics about data collection and modeling are in **Table 1**.

#### Preparation of radiolabeled RNAs

Short target RNAs for binding assays (**Table S1**) were chemically synthesized (IDT) with a 5′ hydroxyl, purified on a 15% urea-polyacrylamide gel, and resuspended in water. They were 5′-radiolabeled using T4 Polynucleotide Kinase (New England Biolabs, M0201) and [γ-^32^P] ATP (PerkinElmer, BLU035C005MC) in T4 PNK reaction buffer (New England Biolabs) at 37°C for 1.5 h, followed by desalting using Micro Bio-Spin P-6 columns (Bio-Rad, 7326221), purification on a 15% urea-polyacrylamide gel, and resuspension in water.

Longer target RNAs for slicing assays were transcribed from single-stranded DNA (IDT) templates (**Table S1**). Template DNAs were purified on a 15% urea-polyacrylamide gel, and annealed with an oligonucleotide containing the T7 promoter sequence (IDT) (**Table S1**). The annealed templates were in vitro-transcribed with in-house purified T7 RNA Polymerase in 5 mM ATP, 2 mM UTP, 5 mM CTP, 8 mM GTP, 5 mM DTT, 40 mM Tris pH 7.9, 2.5 mM spermidine, 26 mM MgCl_2_, 0.01% (v./v.) Triton X-100, 5 mM DTT, SUPERase•In (1 U/µL, Invitrogen, AM2694), and thermostable Inorganic Pyrophosphatase (0.0083 U/µL, New England Biolabs, M0296). After incubation at 37°C for 3−4 h, then a further incubation with RQ1 DNase (Promega, M6101) at 37°C for 30 min, RNAs were purified on a 15% urea-polyacrylamide gel and resuspended in water. To radiolabel, RNAs were dephosphorylated using Quick CIP (New England Biolabs, M0525) at 37°C for 15 min, followed by heat-inactivation at 80°C for 3 min. Dephosphorylated RNAs were radiolabeled using T4 Polynucleotide Kinase and [γ-^32^P] ATP in CutSmart buffer (New England Biolabs) supplemented with 5 mM DTT, at 37°C for 1.5 h, followed by desalting using Micro Bio-Spin P-30 columns (Bio-Rad, 7326250), purification on a 15% urea-polyacrylamide gel, and resuspension in water.

Guide RNAs (**Table S1**) were synthesized with a 5′ monophosphate (IDT), purified on a 15% urea-polyacrylamide gel, and resuspended in water. They were dephosphorylated using shrimp alkaline phosphatase (New England Biolabs, M0371) at 37°C for 30 min, followed by heat-inactivation at 75°C for 5 min, and desalting using Micro Bio-Spin P-6 columns. They were then radiolabeled using T4 Polynucleotide Kinase and [γ-^32^P] ATP in T4 PNK reaction buffer at 37°C for 1.5 h, with a subsequent chase with 0.16 mM ATP at 37°C for 15 min. RNAs were desalted using Micro Bio-Spin P-6 columns, purified on a 15% urea-polyacrylamide gel, and resuspended in water.

#### Small-scale purification of HsAGO2−guide complexes for biochemical assays

A wildtype pcDNA3.3−3xFLAG−SUMO^Eu1^−HsAGO2 plasmid was first generated from the pcDNA3.3−3xFLAG−SUMO^Eu1^−HsAGO2^D669A^ plasmid through site-directed mutagenesis by PCR with KAPA HiFi HotStart Ready Mix. Plasmids with mutant HsAGO2 sequences were then generated by site-directed mutagenesis with appropriate primers (**Table S1**). Plasmids were prepared using the Plasmid Plus Midi Kit (QIAGEN, 12945), and confirmed by Sanger sequencing and whole-plasmid sequencing.

RISCs were purified with a method modified from published protocols.^32^ Guide−passenger RNA duplexes were annealed using 1 µM each of guide and passenger RNAs, with 0−1% ^32^P-radiolabeled guide RNA, in annealing buffer (30 mM Tris-HCl pH 7.5, 100 mM NaCl, and 1 mM EDTA). The mixtures were heated to 90°C and slowly cooled to 30°C over 1.5 h, followed by chilling on ice and storing at −80°C. These duplexes were then used to assemble RISCs in lysates overexpressing the appropriate 3xFLAG−SUMO^Eu1^-tagged HsAGO2 mutants. To generate the lysate, 200 mL of Expi293F cells, cultured in Expi293 expression media at 37°C in 5% CO_2_, were transfected at ∼2 million per mL density, using 190 µg of pcDNA3.3−3xFLAG−SUMO^Eu1^−HsAGO2 (or one of its mutant counterparts), 10 µg pMaxGFP, and 600 µg of polyethylenimine, incubated in 10 mL of Opti-MEM at room temperature for 20 min. After 40 h, cells were harvested and lysed in 8 mL of hypotonic buffer (10 mM HEPES pH 8.0, 10 mM KOAc, 1.5 mM Mg(OAc)_2_, 2% [v./v.] glycerol, 0.5 mM TCEP, 5 mM NDSB-256, with 1 tablet per 50 mL cOmplete EDTA-free Protease Inhibitor Cocktail) by a Dounce homogenizer. Cell lysate was clarified by centrifugation at 3000*g* at 4°C for 15 min, then again for 30 min. The supernatant was re-equilibrated by adding 25% volume of re-equilibration buffer (150 mM HEPES pH 8.0, 700 mM KOAc, 15 mM Mg(OAc)_2_, 42% [v./v.] glycerol, 0.5 mM TCEP, 5 mM NDSB-256, with 1 tablet per 10 mL cOmplete Mini EDTA-free Protease Inhibitor Cocktail), before further clarification by centrifugation at 60,000*g* at 10°C for 20 min. Single-use aliquots were flash-frozen in liquid nitrogen for storage at −150°C.

To assemble each RISC, 900 µL of cell lysate was incubated with 100 µL of 1 µM annealed guide−passenger duplex for 2 h at 25°C. The incubated lysate was centrifuged at 21,000*g* at 4°C for 10 min, and the supernatant was incubated with 150 µL of a slurry of Dynabeads MyOne Streptavidin C1 (Invitrogen, 65002) pre-bound to 75 pmol of capture oligonucleotide (**Table S1**), at 25°C with shaking at 1,300 rpm for 1.25 h. Pre-bound streptavidin beads were prepared by equilibrating the beads according to manufacturer’s instructions, followed by incubation with 75 pmol of 3′-end biotinylated, fully 2′-*O*-methylated oligonucleotides (IDT) containing a seed-only 8mer site for the guide RNA (**Table S1**), at 25°C with shaking at 1,300 rpm for 30 min. The beads were then equilibrated with equilibration buffer (18 mM HEPES pH 7.4, 100 mM KOAc, 1 mM Mg(OAc)_2_, 0.01% [v./v.] IGEPAL CA-630 [Sigma-Aldrich, I3021], 0.1 mg/mL BSA [New England Biolabs, B9000], and 0.01 mg/mL yeast tRNA [Life Technologies, AM7119]) before use. After incubation with lysate, beads were washed three times with 200 µL equilibration buffer and three times with 200 µL capture-wash buffer (18 mM HEPES pH 7.4, 2 M KOAc, 1 mM Mg(OAc)_2_, 0.01% [v./v.] IGEPAL CA-630, 0.1 mg/mL BSA, and 0.01 mg/mL yeast tRNA), then incubated with 112.5 pmol of 3′-biotinylated DNA competitor oligonucleotides (IDT) complementary to the capture oligonucleotides (**Table S1**) in 18 mM HEPES pH 7.4, 1 M KOAc, 1 mM Mg(OAc)_2_, 0.01% (v./v.) IGEPAL CA-630, 0.1 mg/mL BSA, and 0.01 mg/mL yeast tRNA, at 25°C with shaking at 1,300 rpm for 2 h. The eluate was then incubated with 20 µL of anti-FLAG M2 magnetic bead slurry (Millipore, M8823) pre-equilibrated in equilibration buffer according to manufacturer’s instructions, at 25°C with shaking at 1,100 rpm for 2 h. The beads were washed twice with 200 µL of equilibration buffer and twice with 500 µL of equilibration buffer, before incubation in 60 µL of SENP^EuB^ elution solution (500 nM of SENP^EuB^ in equilibration buffer) at 4°C with shaking at 1,300 rpm for 1 h. The eluate was supplemented with glycerol and DTT to a final storage buffer condition of 13.0 mM HEPES pH 7.4, 72.3 mM KOAc, 0.723 mM Mg(OAc)_2_, 5 mM DTT, 0.0723 mg/mL BSA, 0.00723 mg/mL yeast tRNA, 0.00723% (v./v.) IGEPAL CA-630, and 20% (v./v.) glycerol, before flash-freezing in liquid nitrogen for storage at −80°C.

Each purified RISC was run on a 15% urea-polyacrylamide gel alongside the guide−passenger duplex used for its purification. Gels were frozen at −20°C while exposing a phosphorimager plate, and radioactivity was then imaged using the Amersham Typhoon (Cytiva) phosphorimager. Images were quantified using ImageQuant TL (Cytiva) software. Concentration of the RISC was calculated from the radioactivity of the guide in the purified RISC relative to that in the duplex, for which the concentration was known.

RISCs loaded with miR-200b or let-7a were purified without radiolabeling to avoid interfering the signal in dissociation-kinetics assays. Quantification of these complexes was done using a titration assay. A serial dilution of limiting concentrations of RISCs were incubated with either 1 nM (miR-200b RISCs) or 2 nM (let-7a RISCs) of radiolabeled binding-target RNA in 16.5 mM HEPES pH 7.4, 91.7 mM KOAc, 0.917 mM Mg(OAc)_2_, 5 mM DTT, 0.03 mg/mL BSA, 0.003 mg/mL yeast tRNA, 0.01% IGEPAL CA-630, and 6% (v./v.) glycerol, at 37°C for 1 h. To carry out filter-binding assays to quantify RNA-binding, nitrocellulose (Amersham Protran, 0.45 µm pores; Cytiva, 10600062) and nylon (Amersham Hybond-XL; Fisher Scientific, 45001147) membrane filters were cut into discs of 0.5-inch diameter and equilibrated at 25°C for at least 20 min in filter-binding buffer (18 mM HEPES pH 7.4, 100 mM KOAc, 1 mM Mg(OAc)_2_). Each nitrocellulose disc was stacked on top of a nylon disc, then placed on a circular pedestal mounted on a Visiprep SPE Vacuum Manifold (Supelco, 57250-U), set at approximately−20 kPa. 10 µL of reaction was applied to stacked filter membrane discs, followed by 100 µL of ice-cold filter-binding wash buffer (18 mM HEPES pH 7.4, 100 mM KOAc, 1 mM Mg(OAc)_2_, 5 mM DTT). Filter membrane discs were separated, air-dried, then imaged by phosphorimaging and quantified using ImageQuant TL software. Fractions of target RNA bound across RISC dilutions were fit to a quadratic equation by nonlinear least-squares regression in R using the Levenberg-Marquardt algorithm (nlsLM from the R package minpack.lm):

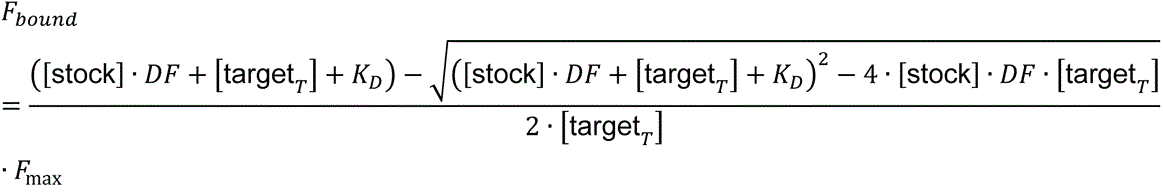

where *F*_bound_ represents fraction of target bound, [target*_T_*] represents the concentration of total target oligonucleotide, [stock] represents stock concentration of RISC, *DF* represents the dilution factor, *K*_D_ represents the dissociation constant for the affinity between RISC and the target, and *F*_max_ represents the maximal fraction of target bound at the plateau. For RISCs programed with miR-200b, [stock] was initialized at 100,000 pM and limited to the numerical range (0, 10^6^); *K*_D_ was initialized at 10^1.5^ pM, limited to the numerical range (10^0.5^, 10^2.5^), and fit in log-transformed space; *F*_max_ was fixed at 1. For RISCs programed with let-7a, [stock] was initialized at 100,000 pM and limited to the numerical range (0, 2×10^6^); *K*_D_ was initialized at 10^1.5^ pM, limited to the numerical range (10^−10^, 10^10^), and fit in log-transformed space; *F*_max_ was initialized at 0.9 and limited to the numerical range (0, 1).

#### Slicing assays

Slicing assays were conducted and analyzed similarly to previously described methods.^32^ Briefly, assays were conducted as single-turnover reactions including 0.05 nM radiolabeled target RNA and a dilution series of RISCs at 2, 5, or 10 nM, in 15.5 mM HEPES pH 7.4, 86.2 mM KOAc, 0.862 mM Mg(OAc)_2_, 5 mM DTT, 0.05 mg/mL BSA, 0.005 mg/mL yeast tRNA, 0.01% (v./v.) IGEPAL CA-630, and 10% (v./v.) glycerol, at 37°C. At time points, aliquots were quenched by mixing rapidly with 1.5X volume of gel-loading buffer at 4°C, then denatured at 90°C for 1 min and run on a 15% urea-polyacrylamide gel. Gels were frozen at −20°C while exposing a phosphorimager plate, and radioactivity was imaged using a phosphorimager and quantified using ImageQuant TL.

Data points were first fit by nonlinear least-squares regression in R using the Levenberg-Marquardt algorithm (nlsLM) to an approximation equation, which assumes pseudo-steady-state of enzyme-substrate complex concentration, to generate guesses for parameter values:

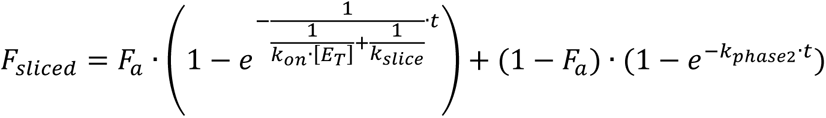

where *F*_sliced_ represents fraction of target sliced, *t* represents time in s, *k*_on_ represents the association rate constant in nM^−1^ s^−1^, [E_T_] represents the total concentration of RISC in nM, *k*_slice_ represents the slicing rate constant in s^−1^, *k*_phase2_ represents the slow second-phase slicing rate constant in s^−1^, and *F*_a_ represents the height of the first phase. *k*_on_ was initialized at 0.02 nM^−1^ s^−1^ and limited to the numerical range (10^−6^, 0.1), where 0.1 nM^−1^ s^−1^ was the diffusion limit. *k*_slice_ was initialized at either 0.0167 s^−1^ or 1.67 × 10^−5^ s^−1^ for very slow reactions, and limited to the numerical range (10^−6^, 1). *k*_phase2_ was initialized at 3.33 × 10^−6^ s^−1^ and limited to the numerical range (1.67 × 10^−6^, 1.67 × 10^−4^), where 1.67 × 10^−6^ s^−1^ was the approximate detection limit based on the longest time points tested. *F*_a_ was initialized at 0.95 × the highest fraction of target sliced measured in the dataset, or at 0.85 for reactions too slow to reach the plateau at the longest time points tested, and limited to the numerical range (0.75, 0.999). If the nonlinear least-squares regression failed, the initial values were used with *F*_a_ adjusted to 0.85.

These guesses were then used as initial values to fit the dataset to an ODE model to calculate accurate values:

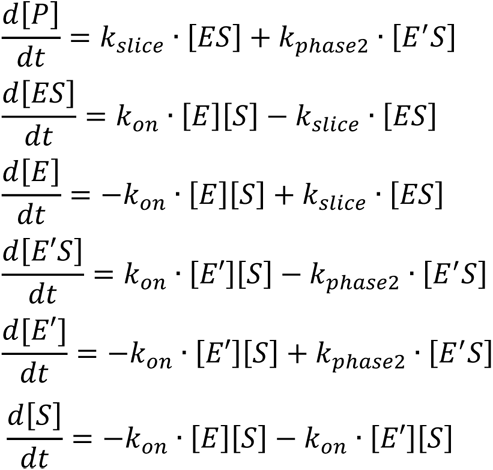

with

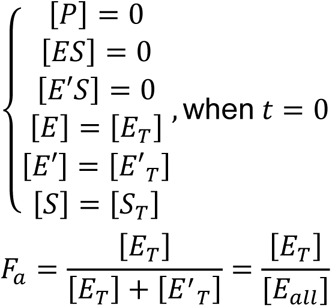

such that, at each time point,

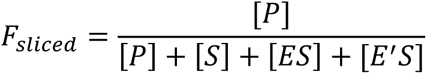

where *F*_sliced_ represents fraction of target sliced, *t* represents time in s, [E_T_] represents total concentration of functionally intact RISC in nM, [E’_T_] represents total concentration of defective RISC in nM, [E_all_] represents total concentration of all RISCs, *F*_a_ represents the fraction of RISCs that is functionally intact, *k*_on_ represents the association rate constant in nM^−1^ s^−1^, *k*_slice_ represents the slicing rate constant in s^−1^, and *k*_phase2_ represents the defective slicing rate constant in s^−1^. Fitting to the ODE model was carried out with the Levenberg-Marquardt algorithm using the modFit function (from the FME package in R), minimizing the total absolute deviation in the predicted fraction sliced at each time point as calculated by the Livermore solver for ordinary differential equations (LSODE) from the deSolve package (ode function) in R, using the backward differentiation formula. Kinetic constants were fit in log-transformed space. *k*_on_ was limited to the numerical range (0, 0.1), where 0.1 nM^−1^ s^−1^ was the diffusion limit. *k*_slice_ was limited to the numerical range (0, ∞). *k*_phase2_ was limited to the numerical range (1.67 × 10^−6^, 3.33 × 10^−3^). *F*_a_ was limited to the numerical range (0.6, 1.0). If *k*_on_ could not be confidently fit for fast-binding reactions due to trivial contribution from binding kinetics, model fitting was repeated with *k*_on_ constrained to the diffusion limit at 0.1 nM^−1^ s^−1^. If *k*_phase2_ could not be confidently fit due to insufficiently long time points to resolve the second phase, model fitting was repeated with *k*_phase2_ constrained to zero. Slicing kinetics of HsAGO2^R97E, K98E^−miR-7 and HsAGO2^H56A, R68A, R97A, K98A^−miR-7 for the 2−16-paired substrate, and all RISCs for position-10/11-mismatched targets, were too slow for the plateau to be resolved even at the longest time points, so fitting was conducted with the value of *F*_a_ constrained to be 0.85.

#### Target dissociation assays

Membrane discs and the vacuum manifold for filter binding were prepared as described above. For each reaction, 6 nM of HsAGO2 (or mutants) loaded with either hsa-miR-200b or hsa-let-7a was incubated with 0.5 nM of the respective radiolabeled binding target, which features seed + supplementary complementarity (**Table S1**), in 23.3 mM HEPES pH 7.4, 129 mM KOAc, 1.30 mM Mg(OAc)_2_, 7 mM DTT, 0.04 mg/mL BSA, 0.004 mg/mL yeast tRNA, 0.014% (v./v.) IGEPAL CA-630, and 8% (v./v.) glycerol, at 37°C for 1 h. The reaction was then diluted in five-fold volume into 100 nM of non-radiolabeled target RNA (1000-fold excess over radiolabeled), with the final reaction condition being 17.6 mM HEPES pH 7.4, 97.8 mM KOAc, 0.978 mM Mg(OAc)_2_, 5 mM DTT, 0.008 mg/mL BSA, 0.0008 mg/mL yeast tRNA, 0.010% (v./v.) IGEPAL CA-630, and 1.6% (v./v.) glycerol, at 37°C. At each time point (determined to the closest second), a 10 µL aliquot was applied to stacked membranes, followed by 100 µL of ice-cold filter-binding wash buffer. Filter membrane discs were separated, air-dried, then imaged by phosphorimaging and quantified using ImageQuant TL software. Fractions of target RNA bound over time points were fit to an exponential equation by nonlinear least-squares regression in R using the Levenberg-Marquardt algorithm (nlsLM from the R package minpack.lm) for no more than 1000 iterations:

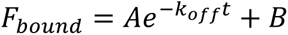

where *F*_bound_ represents the fraction of target RNA bound, *k*_off_ represents the dissociation rate constant in s^−1^, *t* represents time in s, *A* represents the initial bound fraction without background, and *B* represents the background. *k*_off_ was initialized at *e*^0.2^ = 1.22, limited to the numerical range (0, ∞), and fit in log-transformed space. *A* was initialized at 0.75 and limited to the range (0, 1). *B* was initialized at 0.05 and limited to the range (0, 1).

#### Evolutionary analyses of AGO homologs

Peptide sequences were downloaded from UniProt.^70^ Multiple sequence alignment was conducted with the MUSCLE algorithm^71^ using the SnapGene software. Alignment was carried out using 100 representative bilobed AGO- or PIWI-family proteins across the evolutionary tree, including 35 AGOs and 23 PIWIs from 12 animal species ranging from sponges to humans, 22 AGOs from eight plant species spanning *Chlamydomonas reinhardtii* (green algae), *Physcomitrium patens* (bryophyte), and *Arabidopsis thaliana* (eudicot), three AGOs and three PIWIs from four protozoan species, and six fungal AGOs and eight prokaryotic AGOs previously studied in the literature.

#### Major-groove measurements of RNA models

RNA major-groove widths were measured using the web 3DNA server v2.4.3-2019apr06.^72^ Refined inter-phosphorus distances were used, subtracting 5.8 Å to account for the van der Waals radii of phosphate groups.^72,73^ Standard A-form double helices of sequence-matched RNAs, starting at position 2, were generated in ChimeraX for comparison.

