## Supplementary Figures for "The structural basis for RNA slicing by human Argonaute2"

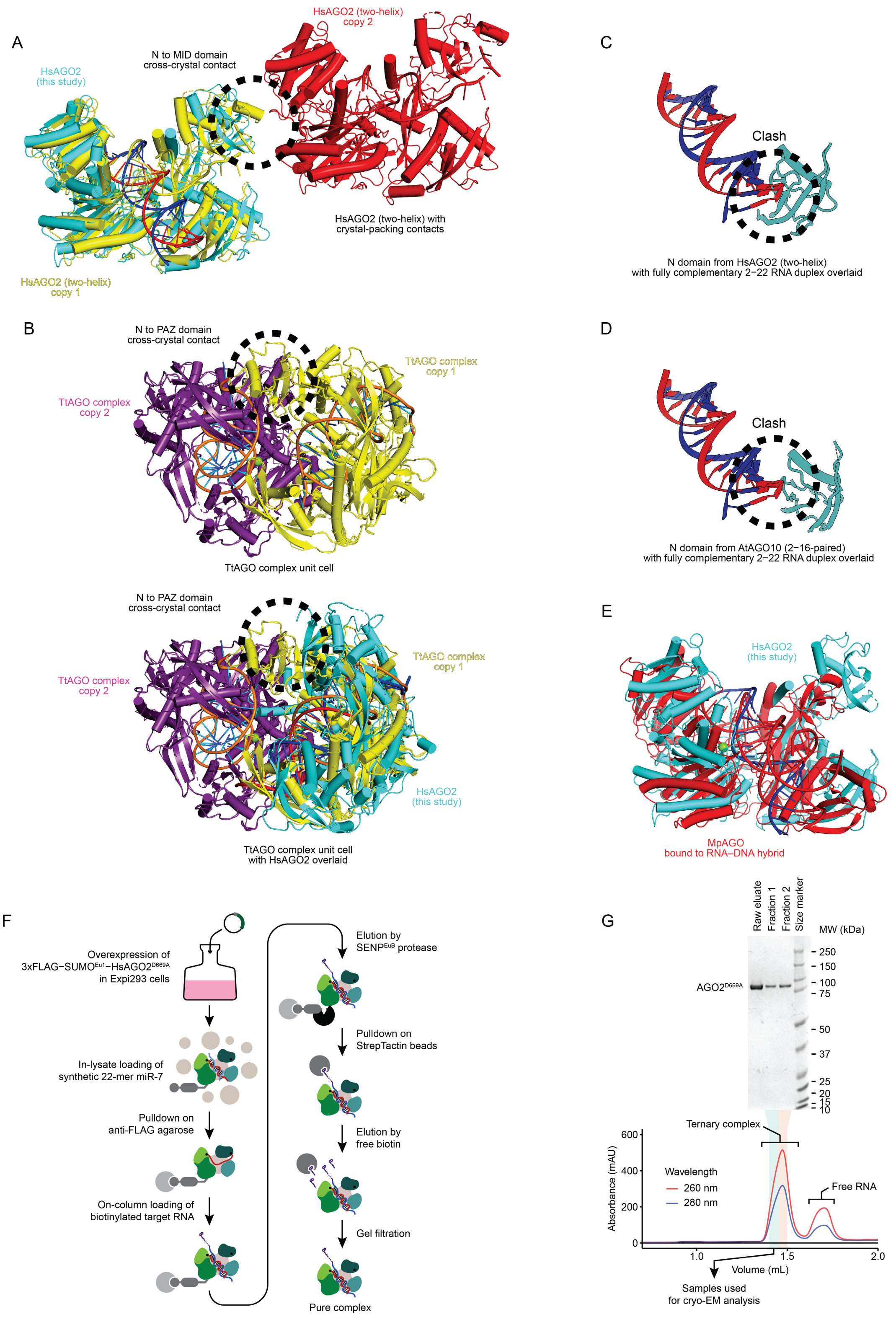

**Figure S1. Structural comparisons and purification of the HsAGO2−miR-7−target ternary complex.**
(A) Crystal-packing contacts for HsAGO2 RISC in the two-helix conformation (PDB: 6N4O[31]). N and MID domains form packing contacts between two copies of HsAGO2 RISC in the two-helix conformation (colored red and yellow). The fully paired HsAGO2 structure (cyan) is overlaid, showing that it is in a conformation that would disrupt these packing interactions.
(B) Crystal contacts for TtAGO complex with a complementary target. At the top is the asymmetric unit of TtAGO, which contains two copies of TtAGO (colored yellow and purple) (PDB: 4NCB[35]), which form contacts between the N and PAZ domains. At the bottom the fully paired HsAGO2 structure (cyan) is overlaid, showing that it would not accommodate the packing contacts.
(C) Clash observed when the N domain from the HsAGO2 two-helix structure (PDB: 6N4O[31]) is overlaid with a 21-bp RNA duplex model generated in ChimeraX bound within the central channel.
(D) Clash observed with the N domain when the RNA duplex from the AtAGO10 2−16-paired structure (PDB: 7SWF[34]) is extended to position 22.
(E) MpAGO bound to an RNA−DNA hybrid fully paired to position 20 (PDB: 5UXO[38]) (red). The fully paired HsAGO2 structure (cyan) is overlaid.
(F) Purification scheme for the $HsAGO2^{D669A}$−miR-7−target ternary complex.
(G) Size-exclusion chromatography of the ternary complex on a Superdex 200 3.1/200 column, followed by analysis on an SDS-polyacrylamide gel, visualizing protein with Imperial (Coomassie R-250) staining.

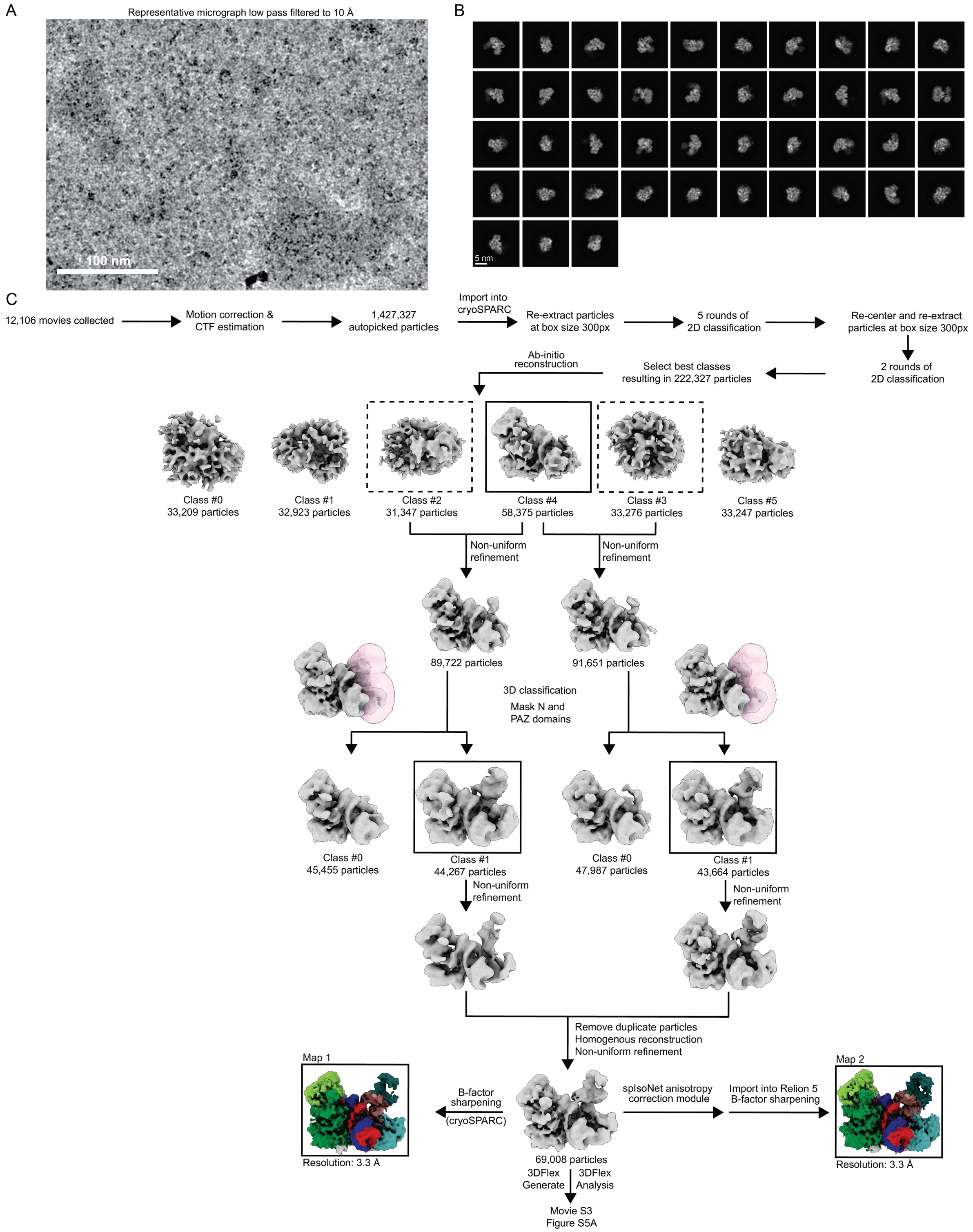
A
Representative micrograph low pass filtered to 10 Å
100 nm
B
5 nm
C
12,106 movies collected
Motion correction & CTF estimation
1,427,327 autopicked particles
Import into cryoSPARC
Re-extract particles at box size 300px
5 rounds of 2D classification
Re-center and re-extract particles at box size 300px
2 rounds of 2D classification
Select best classes resulting in 222,327 particles
Ab-initio reconstruction
Class #0 33,209 particles
Class #1 32,923 particles
Class #2 31,347 particles
Class #4 58,375 particles
Class #3 33,276 particles
Class #5 33,247 particles
Non-uniform refinement
Non-uniform refinement
89,722 particles
91,651 particles
3D classification
Mask N and PAZ domains
Class #0 45,455 particles
Class #1 44,267 particles
Class #0 47,987 particles
Class #1 43,664 particles
Non-uniform refinement
Non-uniform refinement
Remove duplicate particles
Homogenous reconstruction
Non-uniform refinement
Map 1
Resolution: 3.3 Å
B-factor sharpening (cryoSPARC)
69,008 particles
3DFlex Generate
3DFlex Analysis
Movie S3
Figure S5A
spIsoNet anisotropy correction module
Import into Relion 5 B-factor sharpening
Map 2
Resolution: 3.3 Å

**Figure S2. Cryo-EM data collection and processing.**
(A) Representative micrograph low-pass filtered to 10 Å. This micrograph is representative of 12,106 micrographs. Scale bar is shown at the bottom left.
(B) Representative 2D classes. Scale bar is shown.
(C) Classification tree for determining the structure of HsAGO2 in a slicing-competent conformation.

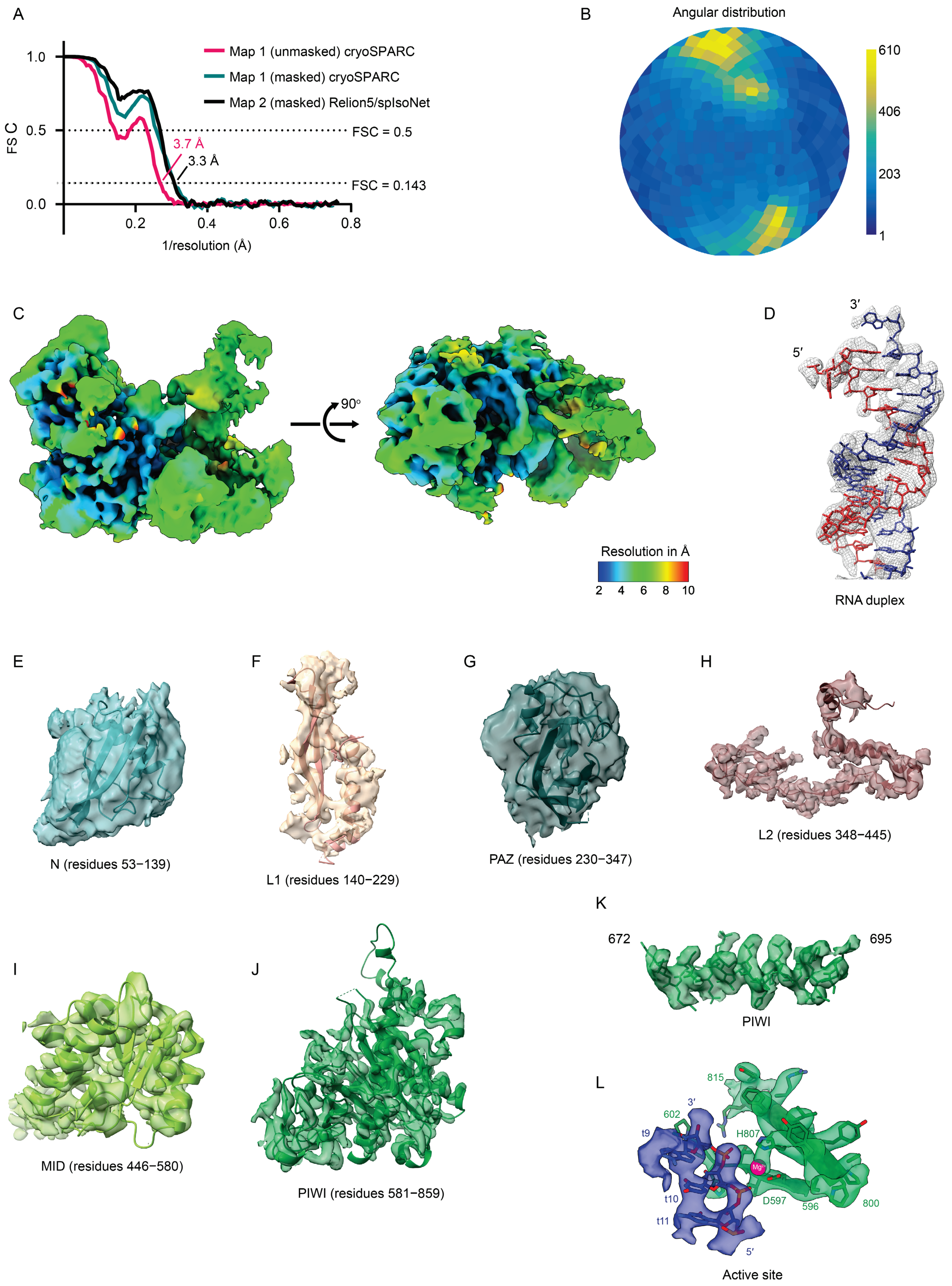
A
Map 1 (unmasked) cryoSPARC
Map 1 (masked) cryoSPARC
Map 2 (masked) Relion5/spIsoNet
FSC = 0.5
FSC = 0.143
3.7 Å
3.3 Å
FSC
1.0
0.5
0.0
0.2
0.4
0.6
0.8
1/resolution (Å)
B
Angular distribution
610
406
203
1
C
90°
Resolution in Å
2 4 6 8 10
D
3′
5′
RNA duplex
E
N (residues 53−139)
F
L1 (residues 140−229)
G
PAZ (residues 230−347)
H
L2 (residues 348−445)
K
672
695
PIWI
I
MID (residues 446−580)
J
PIWI (residues 581−859)
L
815
3′
602
t9
H807
D597
596
800
t10
t11
5′
Active site

**Figure S3. Quality and resolution of cryo-EM data.**
(A) Estimate of average resolution. Dotted lines indicate Fourier shell correlation (FSC) of 0.5 and 0.143. Solid lines indicate FSC between half-maps of the reconstruction.
(B) Angular distribution plot for particles used to reconstruct Map 1. Shading from blue to yellow indicates the number of particles at a given orientation.
(C) Reconstruction of the fully paired HsAGO2 complex, colored by local resolution (Map 1).
(D) Model of the RNA duplex in density from Map 2.
(E) Model of the N domain in density from Map 2.
(F) Model of the L1 domain in density from Map 2.
(G) Model of the PAZ domain in density from Map 2.
(H) Model of the L2 domain in density from Map 2.
(I) Model of the MID domain in density from Map 2.
(J) Model of the PIWI domain in density from Map 2.
(K) Model of PIWI-domain residues 672−695 in density from Map 2.
(L) Model of active site in density from Map 2.

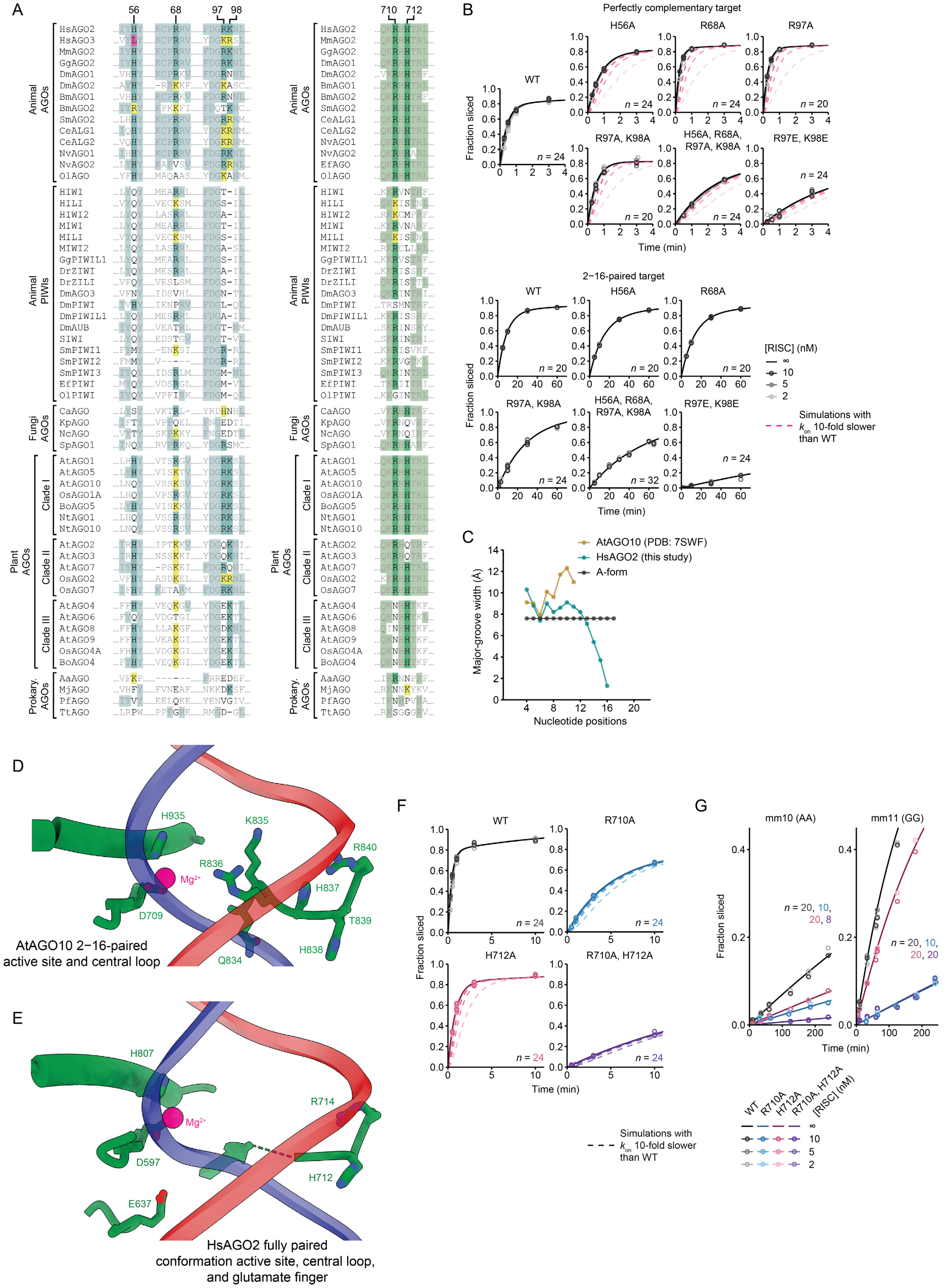

A
56
68
97 98
710 712
Animal AGOs
Animal PIWIs
Fungi AGOs
Plant AGOs
Clade I
Clade II
Clade III
Prokary. AGOs
HsAGO2
HsAGO3
MmAGO2
GgAGO2
DmAGO1
DmAGO2
BmAGO1
BmAGO2
SmAGO2
CeALG1
CeALG2
NvAGO1
NvAGO2
OlAGO
HIWI
HILI
HIWI2
MIWI
MILI
MIWI2
GgPIWIL1
DrZIWI
DrZILI
DmAGO3
DmPIWI
DmPIWIL1
DmAUB
SIWI
SmPIWI1
SmPIWI2
SmPIWI3
EfPIWI
OlPIWI
CaAGO
KpAGO
NcAGO
SpAGO1
AtAGO1
AtAGO5
AtAGO10
OsAGO1A
BoAGO5
NtAGO1
NtAGO10
AtAGO2
AtAGO3
AtAGO7
OsAGO2
OsAGO7
AtAGO4
AtAGO6
AtAGO8
AtAGO9
OsAGO4A
BoAGO4
AaAGO
MjAGO
PfAGO
TtAGO
B
Perfectly complementary target
WT
H56A
R68A
R97A
R97A, K98A
H56A, R68A, R97A, K98A
R97E, K98E
Fraction sliced
Time (min)
2–16-paired target
[RISC] (nM)
∞
10
5
2
Simulations with $k_{on}$ 10-fold slower than WT
C
AtAGO10 (PDB: 7SWF)
HsAGO2 (this study)
A-form
Major-groove width (Å)
Nucleotide positions
D
H935
K835
R840
R836
Mg²⁺
H837
D709
T839
H838
Q834
AtAGO10 2–16-paired active site and central loop
E
H807
Mg²⁺
R714
D597
H712
E637
HsAGO2 fully paired conformation active site, central loop, and glutamate finger
F
WT
R710A
H712A
R710A, H712A
Fraction sliced
Time (min)
Simulations with $k_{on}$ 10-fold slower than WT
G
mm10 (AA)
mm11 (GG)
Fraction sliced
Time (min)
WT
R710A
H712A
R710A, H712A
[RISC] (nM)

**Figure S4. Residues in N and PIWI domains regulate HsAGO2 slicing activity.**
(A) On the left is a multiple sequence alignment of select AGO and PIWI homologs at the proposed N-domain contacts. Residues identical to HsAGO2 are shaded in teal. Residues that changed from HsAGO2 but remained basic are shaded in yellow. An H56L substitution in HsAGO3 is shaded in pink. Species codes are as follows: Hs, *Homo sapiens* (human); Mm, *Mus musculus* (mouse); Gg, *Gallus gallus* (chicken); Dm, *Drosophila melanogaster* (fruit fly); Bm, *Bombyx mori* (silkmoth); Ce, *Caenorhabditis elegans* (nematode); Sm, *Schmidtea mediterranea* (flatworm); Nv, *Nematostella vectensis* (starlet sea anemone); Ef, *Ephydatia fluviatilis* (river sponge); Ol, *Oscarella lobularis* (sea sponge); Dr, *Danio rerio* (zebrafish); Ca, *Candida albicans*; Kp, *Kuyveromyces polysporus*; Nc, *Naumovozyma castellii*; Sp, *Schizosaccharomyces pombe* (fission yeast); At, *Arabidopsis thaliana*; Os, *Oryza sativa* (rice); Bo, *Brassica oleracea* (wild cabbage); Nt, *Nicotiana tabacum* (tobacco); Aa, *Aquifex aeolicus*; Mj, *Methanocaldococcus jannaschii*; Pf, *Pyrococcus furiosus*; Tt, *Thermus thermophilus*. On the right is a multiple sequence alignment of select AGO and PIWI homologs at the central loop. Residues identical to HsAGO2 are shaded in green; otherwise, as in the left panel.
(B) Fraction of target RNA sliced over time by either wildtype (WT) or N-domain mutants of HsAGO2−miR-7, across different RISC concentrations (gray gradient). Solid lines represent best-fit lines from fitting to the ordinary differential equation system. The extrapolated reaction curve at infinite RISC concentration, which represents a reaction rate determined by only $k_{slice}$ and not $k_{on}$, is plotted in black. Magenta dashed lines indicate simulated results that would have occurred if there were a substantial (10-fold) defect in elementary rate constant for target association ($k_{on}$). Number of data points for each set is indicated as *n*. Time points beyond the limits of the *x* axes are not shown.
(C) Analysis of major-groove width deviation from A-form of the RNA duplex in HsAGO2 in the slicing-competent conformation and the AtAGO10 2−16-paired conformation (PDB: 7SWF[34]).
(D) Central loop in the AtAGO10 2−16-paired conformation (PDB: 7SWF[34]). Colors are as in **Figure 4D**.
(E) Glutamate finger, central loop, and active site of HsAGO2 in the fully paired conformation. Otherwise, as in **D**.
(F) Fraction of target RNA sliced over time by either wildtype (WT) or central-loop mutants, across different RISC concentrations. Dashed lines indicate simulated results that would have occurred if there were a substantial (10-fold) defect in elementary rate constant for target association ($k_{on}$). Colors are as in **Figure 4F**; otherwise, as in **B**. Values with WT HsAGO2 are replotted from **B** for reference.
(G) Fraction of centrally mismatched target RNAs sliced over time by either wildtype (WT) or central-loop mutants, across different RISC concentrations; otherwise, as in **F**.

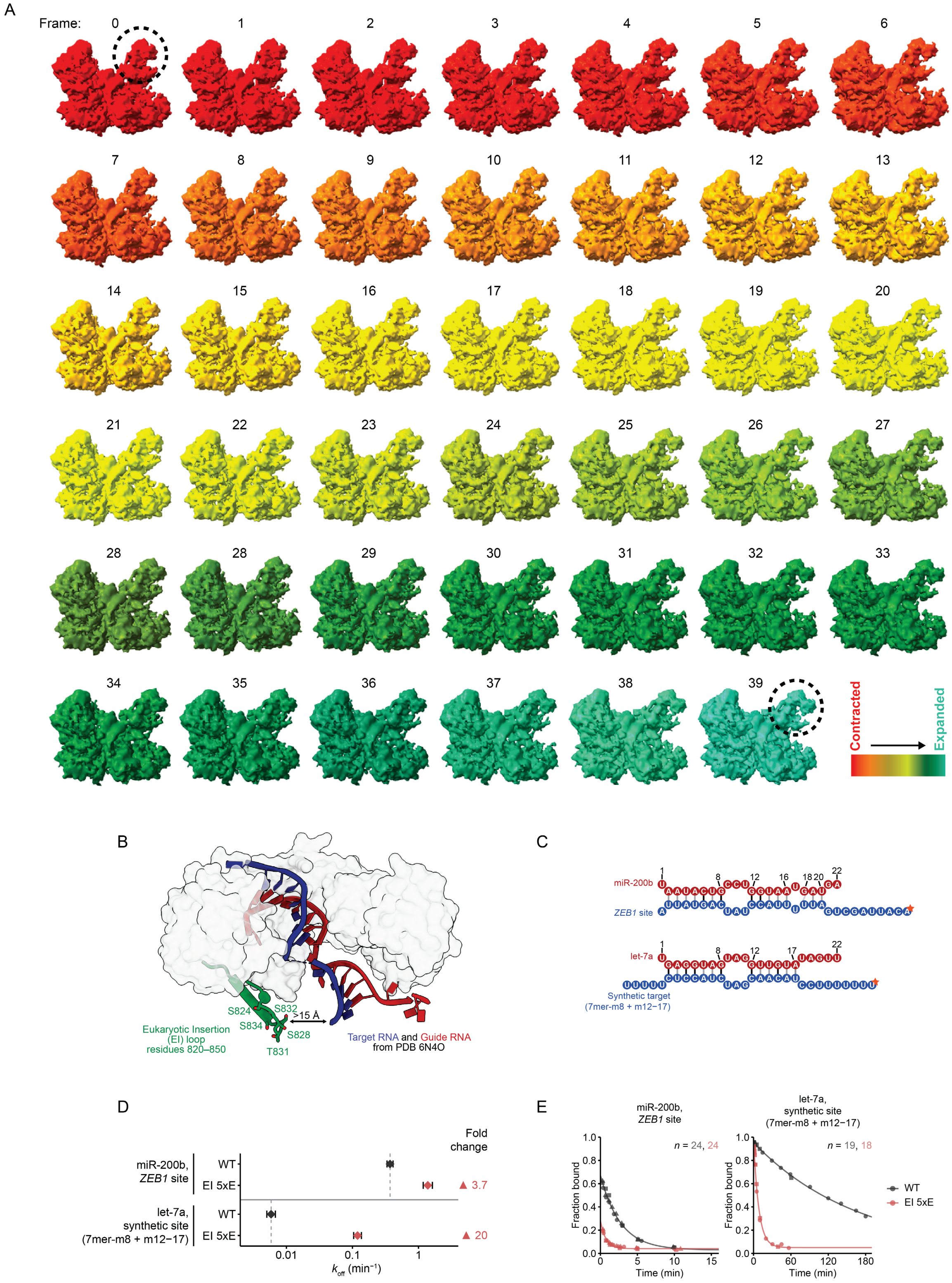

A
Frame:
Contracted
Expanded
B
S824
S832
S834
S828
T831
>15 Å
Eukaryotic Insertion
(EI) loop
residues 820–850
Target RNA and Guide RNA
from PDB 6N4O
C
miR-200b
ZEB1 site
let-7a
Synthetic target
(7mer-m8 + m12–17)
D
Fold
change
miR-200b,
ZEB1 site
WT
EI 5xE
▲ 3.7
let-7a,
synthetic site
(7mer-m8 + m12−17)
WT
EI 5xE
▲ 20
0.01
0.1
1
$k_{off}$ (min⁻¹)
E
miR-200b,
ZEB1 site
n = 24, 24
let-7a,
synthetic site
(7mer-m8 + m12−17)
n = 19, 18
Fraction bound
Time (min)
WT
EI 5xE

**Figure S5. Analyses of the central channel expansion and the EI Loop.**
(A) Frames 0−39 of 3DFlex movie (**Movie S3**). Gradient depicts contracted (red) to expanded (cyan) states.
(B) A ~15 Å distance between the EI loop modeled in our HsAGO2 structure and the target RNA from the HsAGO2 two-helix structure (PDB: 6N4O[31]). The EI loop is shown as a cartoon from residues 820−850, and the rest of HsAGO2 is shown as a surface.
(C) Schematic of miRNA guide–target duplexes examined in target-dissociation assays. The $^{32}P$ radiolabel at the 5′ end of target is indicated as an orange star. Otherwise, this panel is as in **Figure 1B**.
(D) In vitro dissociation-rate constant ($k_{off}$) values for either wildtype (WT) HsAGO2 (dark grey) or HsAGO2 with phosphomimetic substitutions in the EI loop (red), for the two miRNA−target sets tested. Otherwise, this panel is as in **Figure 2E**.
(E) Dissociation of target RNA from either wildtype (WT) HsAGO2 or HsAGO2 with phosphomimetic substitutions in the EI loop. Curves represent nonlinear best-fits to an exponential decay equation. Number of data points for each set is indicated as *n*. Time points beyond the limit of the *x* axes are not shown.
