## Supplementary material for "The structural basis for RNA slicing by human Argonaute2": Table S2

Supplementary Table 2: Modeling of HsAGO2 and nucleic acids

| Domain/ Chain ID | Domain/ region | Residue range | Initial Model | PDB template for initial model/chain | Map used for modeling | Modeling algorithm | Changes to initial model | Confidence of modeling | Relative map resolution |
|---|---|---|---|---|---|---|---|---|---|
| **N-term/A** | N-term | 23-52 | Crystal Structure | 4OLA | 2 | Rigid body fitting | Manual correction, phenix.real_space_refine | Atomic model | 3-8 Å |
| **N/A** | N | 53-140 | Crystal Structure | 4OLA | 2 | Rigid body fitting | Manual correction, phenix.real_space_refine | Atomic model | 6-8 Å |
| **L1/A** | L1 | 141-229 | Crystal Structure | 4OLA | 2 | Rigid body fitting | Manual correction, phenix.real_space_refine | Atomic model | 6-8 Å |
| **PAZ/A** | PAZ | 230-348 | Crystal Structure | 4OLA | 2 | Rigid body fitting | Manual correction, phenix.real_space_refine | Atomic model | 6-8 Å |
| **L2/A** | L2 | 349-444 | Crystal Structure | 4OLA | 2 | Rigid body fitting | Manual correction, phenix.real_space_refine | Atomic model | 3-8 Å |
| **MID/A** | MID | 445-577 | Crystal Structure | 4OLA | 2 | Rigid body fitting | Manual correction, phenix.real_space_refine | Atomic model | 3-8 Å |
| **PIWI/A** | PIWI | 578-821, 845-859 | Crystal Structure | 4OLA | 2 | Rigid body fitting | Manual correction, phenix.real_space_refine | Atomic model | 3-8 Å |
| **PIWI/A** | PIWI/Eukaryotic Insertion Loop | 822-844 | Alphafold2 | N/A | 2 | Alphafold2 | Manual correction, phenix.real_space_refine | Alphafold2 | 6-8 Å |
| **Guide RNA/G** | Guide RNA | 1-8 | Crystal Structure | 6N4O | 2 | Rigid body fitting | Manual correction, phenix.real_space_refine | Atomic model | 3-8 Å |
| **Guide RNA/G** | Guide RNA | 9-21 | ChimeraX generated | N/A | 2 | Rigid body fitting | Manual correction, phenix.real_space_refine | Pseudo-atomic model | 3-8 Å |
| **Target RNA/T** | Target RNA | 15-22 | Crystal Structure | 6N4O | 2 | Rigid body fitting | Manual correction, phenix.real_space_refine | Atomic model | 3-8 Å |
| **Target RNA/T** | Target RNA | 1-14 | ChimeraX generated | N/A | 2 | Rigid body fitting | Manual correction, phenix.real_space_refine | Pseudo-atomic model | 3-8 Å |
